# The ion permeability of DNA nanotube channels

**DOI:** 10.1101/2022.03.04.482952

**Authors:** Naresh Niranjan Dhanasekar, Yi Li, Rebecca Schulman

## Abstract

Techniques from structural DNA nanotechnology make it possible to assemble complex 3-dimensional nanostructures with virtually arbitrary control over their sizes, shapes and features at length scales of 3–100 nm, providing a flexible means for constructing nanoscale devices and machines. Here, we assemble micron-length DNA nanotubes and assess their performance as pipes for controlled ion transport. DNA nanotubes grow *via* assembly of DNA tiles from a seed pore, a 12-helix DNA origami cylinder functionalized with cholesterol, to form a DNA nanotube channel. The central channel of a nanotube can be obstructed via Watson-Crick hybridization of a channel cap, a second DNA origami structure, to the end of a nanotube channel or a nanotube seed pore. Single-channel electrophysiological characterization shows that both nanotube seed pores and nanotube channels display ohmic ion conductance consistent with their central channels’ diameters. Binding of the channel cap reduces the conductances of both DNA nanotube channels and seed pores, demonstrating control of ion-transport through these micron-length channels. Because these channels could be assembled into branched architectures or routed between specific molecular terminals, these results suggest a route to self-assembling nanofluidic devices and circuits in which transport can be controlled using dynamic biomolecular interactions.

## INTRODUCTION

Transport between membrane-bound compartments is a fundamental mechanism of control in biology. Nanoscale channels regulate ion or small molecule flux across membranes^1–3^ and mediate transport of these species between cells.^4^ Such cell-to-cell communication plays important roles in growth and development and is important in the transmission of pathogens such as human immunodeficiency virus (HIV),^5^ herpes simplex virus (HSV),^6^ and prions.^7^

Biological nanopores and channels and synthetically designed nanochannels likewise underpin a range of modern technologies. Third-generation DNA sequencing,^8,9^ protein and enzyme detection,^10–14^ diagnostics, bioelectronics,^15,16^ and synthetic tissues from synthetic compartments in which pores mediate transport^17,18^ all employ these channels. While these nanopores can mediate transport between abutting membranes, longer nanochannels could mediate transport between more distant compartments or endpoints. Such longer nanochannels are often observed in living systems, *e.g*., tunneling nanotubes between cells.^19,20^ Longer nanochannels could mediate the rapid transport of ions or molecules over longer distances by confining them within one-dimensional channels. As such, the ability to synthesize and control transport through longer nanochannels could create opportunities for engineering transport between synthetic devices and different cells in a 3-dimensional culture or between cells in a 3-dimensional tissue and synthetic devices, or could mediate complex patterns of transport between different devices in a system to create nanofluidic circuits.

The nanometer scale of channels suggests that self-assembly is a practical means for their fabrication. Self-assembled biomolecular channels might also be functionalized or grown in such a way that they could be integrated into cell culture or organized in 3-dimensional environments.^21^ While self-assembled membrane proteins have been adopted for use as membrane pores in synthetic devices, methods from DNA nanotechnology make it possible to emulate the structural and functional aspects of naturally occurring membrane channels using easy-to-design DNA nanostructures. Such DNA channels can enter into lipid bilayers and transport currents ranging from 100 to a few 1000’s of picoamperes across these bilayers.^22–31^ DNA channels can act as biosensors,^22,25,27^ cytotoxic agents,^32^ and drug delivery vehicles,^28,33–36^ or organize enzymes to direct a reaction cascade.^37^ In contrast to biological channels, DNA nanopores have dimensions that can be precisely defined and can be site-specifically functionalized using DNA conjugates within their design.^22–27^ A broad array of micron-length channels whose dimensions and structure can be precisely designed and have also been assembled using DNA.^38^

Here we design, self-assemble, and characterize synthetic DNA nanochannels with sub-10 nm diameter and lengths of 2-3 microns and study their capacity to serve as channels for transporting ions. The structures that we study can present specific chemistry at either end to allow, for example, insertion of the channel end into a membrane. They have a radius and exterior structure that can be precisely tuned and can grow to connect two end points connected by molecular labels,^39^ suggesting how these structures might serve as primitives for directing the self-assembly of nanochannels with precisely tuned properties at desired locations. These structures can also self-heal,^40^ suggesting they have capacities for a longer lifetime in environments such as cell medium.

To serve as conduits for transport, such micron-length nanochannels must mediate transport both across lipid membranes and through the extent of the channel. Here we study the flux of ions through these channels across a membrane and measure the extent to which ions travel along the length of these channel *vs*. being lost into bulk through channel walls.

We construct channels using a folded DNA origami structure as a seed or template (Figure 1a^38^ see Supplementary Note 1, Supplementary Figures 1 and 2 for sequences) from which microns-long nanotubes grow via the polymerization of DNA tiles (Figure 1b^38^ see Supplementary Note 2 and Supplementary Figure 3 for sequences). Hydrophobic moieties on the seed can direct the insertion of a seed or seeded nanotube into a membrane to form a transmembrane pore (Figure 1a and b). To measure the relative rates of ion transport across the channel *vs*. through channel walls, we designed and synthesize a self-assembled channel cap (Figure 1c, see Supplementary Note 3 for sequences)^41^ which sharply constricts the channel because its staples directly connect helices on opposite sides of its barrel. We show that these microns-long DNA nanotube channels can form transmembrane pores in lipid bilayers with well-defined conductances that are independent of channel length. Capped seeds and seeded nanotubes (Figure 1c and d) have conductances on order half of their open counterparts suggesting that ions travel both through the channel’s end and across channel walls when a transmembrane potential is applied. This result is consistent with the fact that the walls of DNA nanostructures have crossovers 2 full turns apart, which produces gaps of >1 nm because of electrostatic repulsion.^42,43^

**Figure 1.**
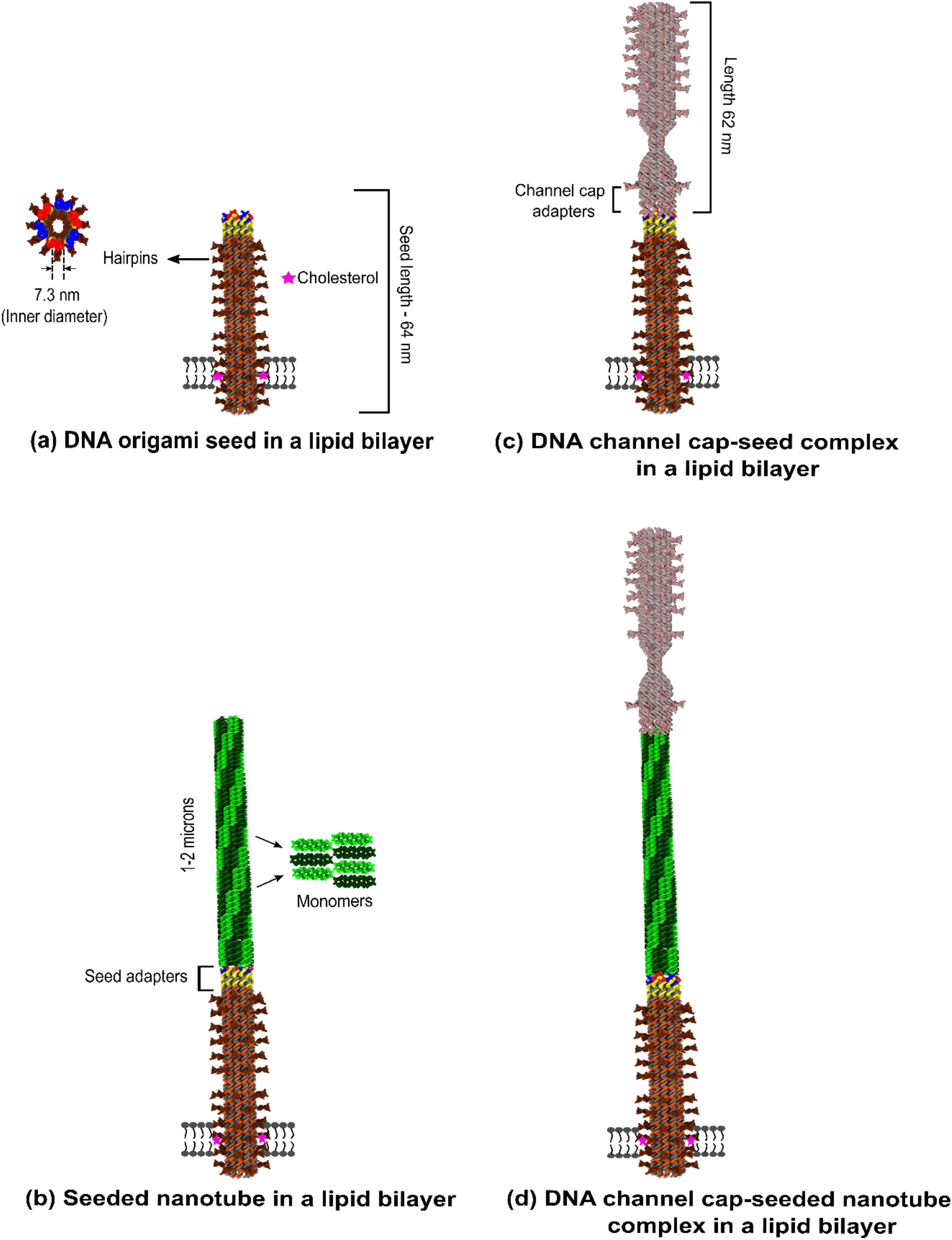
Schematic representations of a DNA origami seed channel, seeded nanotube channel, capped seed channel and capped nanotube channel. (a) The DNA origami seed consists of 72 ssDNA staple strands that fold a long M13mp18 ssDNA into a cylindrical shape. The DNA origami seed has length 64.70 ± 2.12 nm (95% CI, N = 50) with outer and inner diameters of 11 and 7.3 nm, respectively. DNA hairpins on the outer region of the DNA origami seed are designed to direct the seed to cyclize in a specific orientation. The DNA seed is decorated with 18 cholesterol-DNA conjugates. The cholesterol groups are desired to stabilize the structure within a lipid bilayer (For sequences see Supplementary Figures 1 and 2, Supplementary Notes 1 and 4). (b) Nanotubes are grown from two types of monomers (dark green and light green) with different sticky ends. The sticky ends hybridize to form a lattice which in turn cyclizes to form a nanotube which can bind to (or grow from) the nanotube seed (For sequences see Supplementary Figure 3 and Supplementary Note 2). A seeded DNA nanotube in a 1,2-diphytanoyl-*sn*-glycero-3-phosphocholine (DPhPC) lipid bilayer. (c) Hybridization of a DNA seed and a channel cap produces a channel cap-DNA seed complex. A channel cap-DNA seed in a synthetic lipid bilayer is shown (For sequences see Supplementary Notes 3 and 4).^39^ (d) The facet of a seeded tube can hybridize to a channel cap to produce a channel cap-seeded tube complex. A channel cap-seeded DNA tube is shown in a synthetic lipid bilayer.

This study therefore suggests how microns-long DNA nanochannels can mediate ion transport over their length. The assay we develop with a pore and cap could further be used to determine how functionalized or otherwise modified nanotubes might serve as channels through which ions can flow from one channel end to the other. Micron-length channels could be used for sensing, by acting as transport^22,25,27^ conduits with long dwell times, to construct electric circuits,^44^ or for nanofluidic transport.^45,46^

## RESULTS AND DISCUSSION

### DNA origami barrel/nanotube seed channel design

To characterize how microns-long DNA nanostructures might mediate ion transport, we sought to develop a system for assembling DNA nanostructure conduits such that they could mediate transport across a lipid bilayer membrane. We began the design of this structure with a DNA origami seed structure we developed previously.^38^ This structure is a scaffolded DNA origami^47^ consisting of 12 DNA helices arranged in a cylinder (Figure 1a) and whose size has been measured using atomic force microscopy at 63 nm in length and 21 nm in width.^38^ The seed presents single-stranded sticky ends that serve as templates for the assembly of DNA nanotubes that can extend many microns in length (Figure 1b). Seeded growth from these templates provides a means to create channels microns in length. The structures were labeled with fluorophores by binding DNA-fluorophore (ATTO488 or ATTO647N) conjugates to regions of the M13mp18 scaffold not involved in seed formation (see Supplementary Note 4 for sequences).^39^

To create a pore-forming seed structure, we modified a subset of the seed’s staples by adding sequence domains that hybridize to sequences on DNA-cholesterol conjugates (see Supplementary Note 1 for sequences), creating a domain that could enter into a membrane, forming a pore (Figure 1a).^22–24,25–28,30,31^

The average length of nanotube seeds with these modified staples was 64.70 ± 2.12 nm (95% CI, N = 50) (see Materials and Methods), similar to the length of the unmodified structure (Figure 2a and Supplementary Figure 4a, b). Transmission electron micrographs of nanotube seeds with attached DNA-cholesterol conjugates, *i.e*., nanotube seed channels, did not show significant aggregation after assembly, which might be induced by interaction between the cholesterol groups on the conjugates (Figure 2b and Supplementary Figure 4c, d). They also remained dispersed in electrophysiology buffer (1 M KCl, 10 mM MgCl_2_, 10 mM MES) (Figure 2c and Supplementary Figure 4e, f), suggesting that single pores could be studied in electrophysiology experiments. When nanotube seed channels were mixed with small unilamellar vesicles (SUV’s), their barrels spanned the lipid membrane and were consistently oriented perpendicular to it, suggesting that the seeds formed membrane pores (see Materials and Methods, Figure 2d and Supplementary Figure 5).

**Figure 2.**
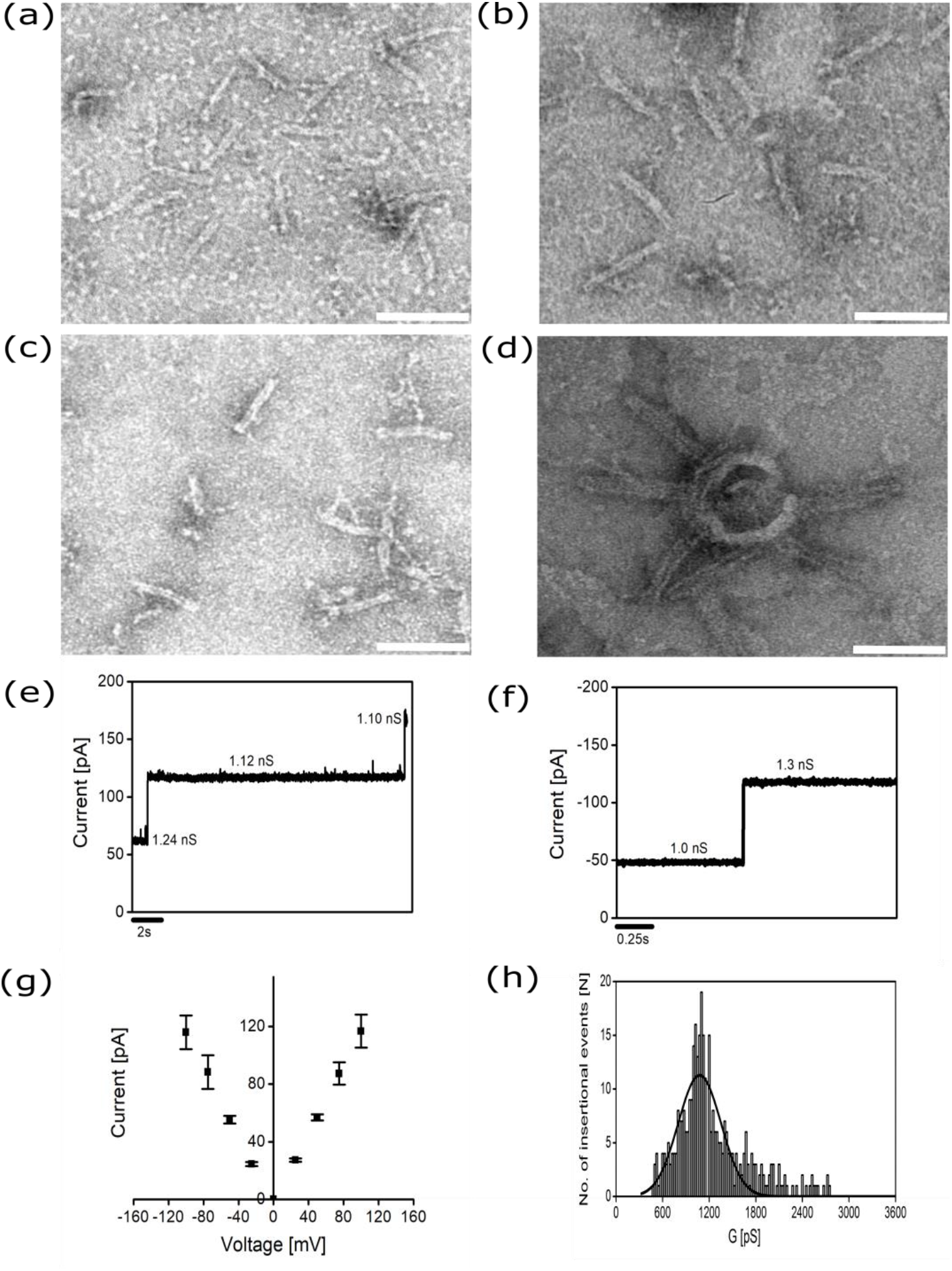
Structure and conductance of DNA origami barrel/nanotube seed membrane channels. (a–d) Transmission electron micrographs of DNA nanotube seed channels. Average channel length is 64.70 ± 2.12 nm (95% CI, N = 50 structures). Scale bars:(a–d) 100 nm. (a) Nanotube seeds in TAEM buffer after filter purification. (b) Nanotube seed channels in TAEM buffer after the addition of cholesterol-DNA conjugates and subsequent filter purification. (c) Nanotube seed channels in electrophysiology buffer (1 M KCl, 10 mM MgCl_2_ buffered with 10 mM 4-morpholine ethane sulfonic acid (MES) monohydrate, pH 6.0). (d) Nanotube seed channels inserted into small unilamellar vesicles (SUVs). Electrical signatures of the DNA origami seed channels in lipid bilayers. (e–f) Insertions of DNA nanotube seed channels in a DPhPC bilayer formed by the droplet interface bilayer (DIB) technique at an applied transmembrane potential of (e) +50 mV. (f)–50 mV. For clarity, here and elsewhere electrical traces were filtered at 200 Hz. (g) Current-voltage (I-V) relationship of the DNA origami nanotube seed channels (N = 3). (h) Histogram of the conductance steps of the nanotube seed channels inserted in DPhPC bilayers formed by the DIB technique at an applied transmembrane potential of +50 mV (N = 362). A Gaussian fit to the conductances is shown as an overlay (1200 ± 275 pS, ±S.D).

### Electrical recordings of the DNA nanostructures

We next sought to measure ion-conductance through the nanotube seed channels by making electrophysiological measurements using the droplet-interface bilayer (DIB) technique.^25,48–50^ Nanotube seed channels were incubated with DPhPC (1,2-diphytanoyl-*sn*-glycero-3-phosphocholine) lipid vesicles for 10 min, after which the insertion of nanotube seed channels into the membrane was triggered by applying a transmembrane potential of +50 mV (see Materials and Methods). The electrical signature was a stable current of 60 pA at +50 mV applied potential, which corresponds to a conductance of 1.2 nS across a nanotube seed channel (Supplementary Figure 6a). Applying a negative transmembrane potential of –50 mV induced a current change of the same magnitude in the opposite direction (Supplementary Figure 6b and c). Multiple stochastic insertions of the nanotube seed channels into DPhPC bilayers produced changes in the current of roughly the same values (Figure 2e and f). As the two droplets forming a bilayer were separated from one another, we observed stepwise reductions in conductance of similar magnitude to the increases observed during insertion, *i.e*., 6 channel closures yielding a total conductance of 5.9 nS (Supplementary Figure 7). Current changes induced by a range of transmembrane voltages were proportional to the transmembrane voltage, indicating that the nanotube seed channels have ohmic conductance (Figure 2g). This is similar to what has been observed in other cholesterol-based,^22,26,27,31^ porphyrin,^24,30^ tocopherol,^25^ ethane-phosphorothiates^23^ and biotin-streptavidin^25^ anchored DNA origami nanochannels.

Interestingly, we did not observe gating events (repeated switching between open and closed states), which have been previously observed for some DNA origami channels. Such gating events are more prevalent in experiments carried out using planar lipid bilayer where variations in membrane pressure are larger than in experiments using the DIB technique employed here^22,31^. A histogram of the measured channel conductances at +50 mV showed a broad distribution of channel conductances with a peak conductance at 1.2 nS with a standard deviation of a Gaussian fit of 0.27 nS (Figure 2h). Occasionally, we observed higher multiples of the single seed channel conductances such as 2.12 and 3.0 nS but the frequency of those events was low, suggesting that these events might be due to simultaneous insertion of two (2.12 nS) or three (3.0 nS) channels, respectively into the lipid bilayer (Supplementary Figure 8). As expected, in the absence of DNA-cholesterol conjugates, no current changes were observed when a transmembrane potential was applied across lipid bilayers in the presence of nanotube seeds (Supplementary Figure 9a). Analogously, no current changes were observed in the presence of 10 μM cholesterol-DNA conjugates (but the absence of nanotube seeds) (Supplementary Figure 9b).

The measured conductance of the nanotube seed channels is lower than would be predicted theoretically by modeling the pore as a nanocylinder with electrically non-permeable walls where the conductance is controlled by the flux of solvent (4.8 nS, Supplementary Note 12).^51^ While the assumptions of this model apply to many biological protein channels, they are less relevant for DNA channels, whose walls are likely to be electronically permeable as they are composed of hydrophilic, negatively changed DNA and have gaps large enough that water could pass through them.^51^ The assumption that ions move at a constant rate through the channel is also unlikely to be satisfied by negatively charged DNA origami nanopores.^52^ As a result, the predictions of this model have both over- and underestimated the conductance of other DNA origami channels.^23,27,30,31^

Having shown that DNA nanotube seeds could insert into the membrane to create pores with well-defined conductance, we next asked whether micron length-DNA nanotubes could also act as membrane channels (Figure 1b). DNA nanotubes were grown from seeds by first filter-purifying the seeds and then adding monomers for assembly over a 34-h incubation period (see Materials and Methods, Supplementary Notes 6 and 7). To make it possible to characterize the yield and lengths of nanotubes, monomers were labeled with a small molecule fluorophore Cy3 on their central strands (see Supplementary Note 2 and Supplementary Figure 3 for sequences). DNA-cholesterol conjugates were added after nanotubes were grown and the resulting structures were filter purified again to remove excess strands from the solution of nanotube membrane channels (see Materials and Methods, Figure 3a, b, Supplementary Note 7 and Supplementary Figure 10).

**Figure 3.**
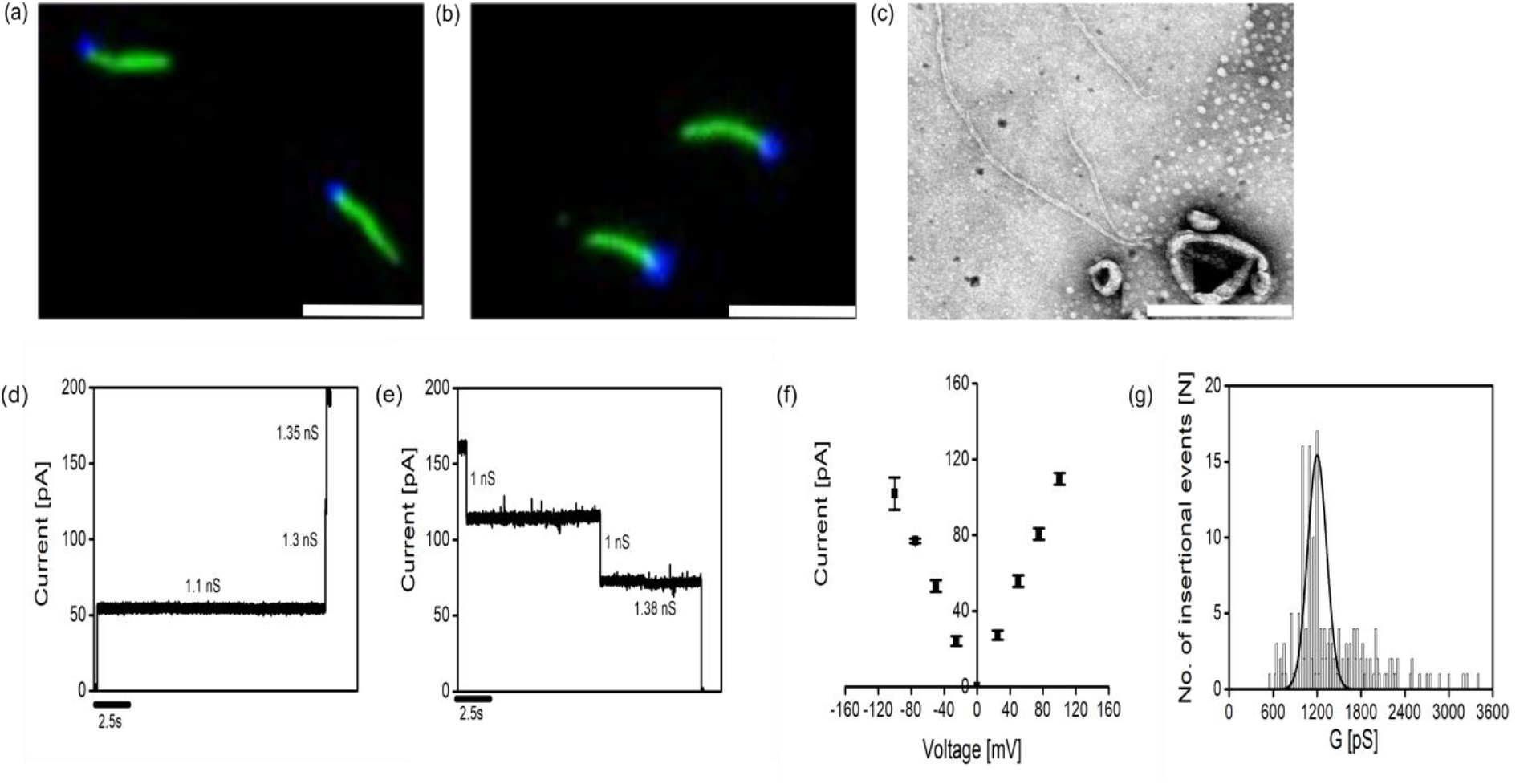
Structure and conductance of DNA seeded nanotube membrane channels. (a–b) Fluorescence micrographs of seeded DNA nanotubes. Nanotubes are labeled with Cy3 (green), nanotube seeds with ATTO488 (blue). Scale bars: 5 μm. (a) Seeded DNA nanotubes after assembly. (b) Seeded nanotube channels after assembly, addition of cholesterol-DNA conjugates and purification. (c) Transmission electron micrograph of seeded DNA nanotube channels in SUV’s. Scale bar: 100 nm. (d) Multiple insertions of DNA seeded nanotube channels inserted into DPhPC bilayers at an applied transmembrane potential of +50 mV. (e) Current decreases observed upon separation of the two droplets forming the lipid bilayer at an applied transmembrane potential of +50 mV. (f) The current-voltage relationship (I-V) observed for single seeded DNA nanotube channels (N = 3). (g) Histogram of the conductance steps of seeded DNA nanotube channels across DPhPC bilayers formed by the DIB at an applied transmembrane potential of +50 mV (N = 186). A Gaussian fit to the conductances is shown as an overlay (1155 ± 128 pS, ±SD).

Fluorescence micrographs showed that 84 ± 3% (95% CI) (N = 678) of DNA nanotube seed pores had nanotubes on them, which was the same as the fraction just after the DNA-cholesterol conjugates were added, 84 ± 1% (95% CI) (N = 467) (Supplementary Figure 11a). The average length of the nanotube channels was 2.43 ± 0.10 μm (95% CI) (N = 185) before purification and 2.17 ± 0.10 μm (95% CI) (N = 272) after purification (Supplementary Figure 11b), suggesting that long nanotubes remained intact after filter purification.

DNA nanotube channels were designed to insert into membranes at the seed ends of the structure, with the channels extending microns away from the anchoring membrane. Transmission electron micrographs confirmed that nanotubes oriented on membranes with the same relative orientation as DNA nanotube seed pores when inserted into DPhPC SUV’s (Figure 3c and Supplementary Figure 12). The curvatures of the nanotubes appeared qualitatively consistent with the previously measured persistence lengths of DNA nanotubes of 8.7 μm.^39^

The conductances of the nanotube channels were measured across a membrane formed by the DIB technique. Nanotube channels had an average conductance of 1.0 nS at an applied transmembrane potential of +50 mV, with different insertions producing different conductances clustered around this value (Figure 3d, e and Supplementary Figures 13 and 14). Control experiments with DNA nanotubes without the seeds produced no significant rises in the current (see Materials and Methods, Supplementary Note 8 and Supplementary Figures 15 and 16). Like nanotube seed channels, the ion conductance of nanotube channels was also ohmic (Figure 3f).

The range of conductances of the nanotubes channels was similar to the range of conductances of nanotube seeds (Figure 3g, Gaussian fit, 1.15 ± 0.12 nS). This range represents the conductance of nanotube channels of different lengths as well as a smaller fraction of seeds without nanotubes, suggesting that the growth of a nanotube from the seed does not change the conductance of the nanotube seed channel. The electron micrographs of SUV’s and nanotubes showed that >500 nm-long seeded nanotubes incorporated into membranes, suggesting that these measurements likely included seeded nanotubes among the measured pores (Figure 3c and Supplementary Figure 12). Thus, DNA nanotubes of various lengths have a single average conductance in the range of 1.0–1.27 nS, similar to the conductance of DNA seed pores. Carbon nanotubes have also been reported to have conductances independent of length: nanotubes of lengths between 5 and 15 nm showed a unitary channel conductance of 630 ± 120 pS,^53^ although here the range of channel lengths is far greater.

We next asked whether we might learn about where ions flow through the seed and nanotube channels to induce conductance across the membrane. Ions could either flow from bulk solution into the opening of the channel of a nanotube seed or nanotube and then across the membrane, or across the DNA walls of the nanotubes or seeds into the channel and then across the membrane. The large gaps between helices in DNA origami structures, which can be created because of electrostatic repulsion of DNA’s negatively charged backbones, could allow ion transport through walls. Leakage of ion-current has been observed in molecular dynamics simulations of other DNA nanopores. ^26,54–57^

If micron-length channels were used to mediate ion transport across multiple membranes, for example (Figure 1c), ion leakage through channel walls in such a long channel would sharply limit the efficiency of this conductivity. To measure the amount of ion-conductance through the channel from end-to-end *vs*. through the channel walls of the DNA seeds and the nanotubes, we constructed a channel cap to attach to the end of a channel (Figures 1 and 4a and see Supplementary Note 3 for sequences) that can bind to and block the channel’s open end. To create the channel cap, we modified the design of a cylindrical nanotube seed that could bind to the end of a DNA nanotube^41^ by adding staples that pulled together the opposite walls of the cylinder, inducing a twist in the structure and constricting its opening (Figures 1 and 4a).^42^ We designed the channel cap to present sticky ends that can bind to the end of either a nanotube or a seed via DNA hybridization to create a stable bond. The gaps between the channel cap and seed or nanotube channel would be expected to be on order of those in the channel walls. We hypothesized that the channel cap, because it reduces the diameter of the channel’s opening, should reduce the ability of ions to cross the membrane because fewer ions can enter the channel at its end and transverse the channel’s length to cross the membrane. The channel cap should not, in contrast, significantly change the rate of ion transport across the DNA walls of the channel. That is, ion flux across a capped nanotube seed or nanotube channel should occur primarily via transport through channel walls (Figure 1).

**Figure 4.**
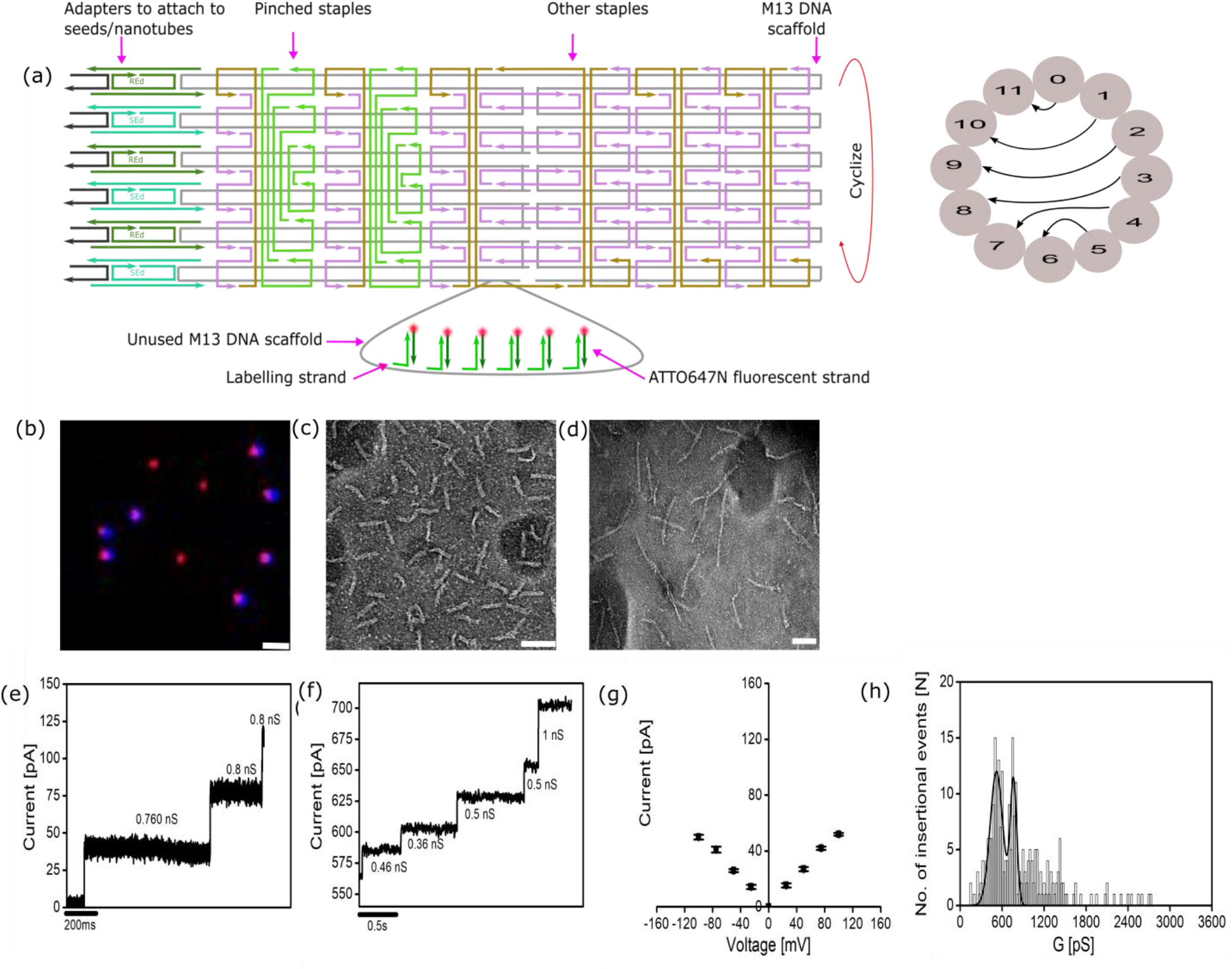
Design and conductivity of channel cap-DNA nanotube seed channels. (a) (Left) Staple strand map (from CadNano) showing the design of the DNA origami channel cap. The sticky ends presented by the adapter strands attach either to the free ends of seeds or to the seeded nanotubes via Watson-Crick hybridization. Pinched staples are those that were modified to reduce the size of the channel opening. (Right) Map of helices showing how they are connected by pinched staples to reduce the opening size. (b) Fluorescence micrographs of seed pores (with attached cholesterol DNA conjugates) with attached channel caps after 2 rounds of filter purification at 3000 g for 30 min. Seeds are labeled with ATTO488 (blue) and channel caps are labeled with ATTO647N (red). Scale bar: 2 μm. (c) Transmission electron micrograph of DNA origami channel caps. Caps have an average length of 62.43 ± 1.21 nm (95% CI, N = 45 structures). Scale bar: 100 nm. (d) Transmission electron micrograph of nanotube seeds bound to channel caps. Average length of the seed-channel cap structure is 132.3 ± 3.2 nm (95% CI, N = 30 structures). Scale bar: 100 nm. (e–f) Multiple insertions of the capped nanotube seed channel in a DPhPC bilayer at an applied transmembrane potential of +50 mV. (g) Current-voltage relationship (I-V) of a capped DNA seed channel (N = 4). (h) Histogram of the conductance steps of capped DNA seed channels across DPhPC bilayers formed by the DIB technique at an applied transmembrane potential of +50 mV (N = 259). A Gaussian fit to the conductances is shown as an overlay (518 ± 90 pS and 762 ± 46 pS).

Channel caps were prepared in a manner similar to the nanotube seeds, *i.e*., using analogous concentrations of staples, the same method of filter purification and the same annealing schedule (see Materials and Methods and Supplementary Note 9). Transmission electron micrographs showed that the channel caps folded into cylinders similar in shape to nanotube seeds and had an average length of 62.43 ± 1.21 nm (95% CI, N = 45 structures). Electron micrographs showed a constriction on one end that also may have introduced a twist (Figure 4c and Supplementary Figure 17).

To verify that channel caps bound to seeds, we incubated nanotube seed channels with channel caps for 3 h at 32°C to allow the channel caps and seeds to bind (see Materials and Methods and Supplementary Note 9) and then prepared the resulting samples for electron microscopy.^41^ The nanotube seeds have adapter strands that bind to one nanotube facet, while the channel caps have adapter strands that bind to a nanotube’s other facet or to the seed. The six adapter sticky end strands on the channel caps and on the nanotube seed should facilitate strong binding of the channel cap to the nanotube seed when the two structures are incubated at 32°C (For complete sequences of adapter strands see Mohammed *et al*).^39^ Consistent with this design, the electron micrographs revealed that the designed channel caps bound well to the nanotube seeds. The average total length of the bound seed-cap structures was 132.3 ± 3.2 nm (95% CI, N = 30 structures) (Figure 4b, d and Supplementary Figure 18). Fluorescence micrographs showed that 91 ± 1% (95% CI, N = 230) of the seeds were bound to channel caps after this incubation (Figure 4b and Supplementary Figures 19 and 20).

To characterize the conductance across the nanotube seed pores whose ends were closed off by channel caps, the channel cap-seed pore complexes were incubated with DNA-cholesterol conjugates (final concentration 10 μM) for 1 h under ambient conditions and were then filter purified in the same manner as nanotube seed pores. After purification, the fraction of seed channels bound to channel caps was unchanged from before purification: 90 ± 1% (95% CI, N = 223) (Supplementary Figures 19 and 20).

The histogram of conductances induced by the nanotube seed channels with attached channel caps when inserted into the membrane had two ranges of conductance peaks: one between 428 and 608—around half the potential of the nanotube seed channels— and the other between 716 and 808 pS at a positive transmembrane potential of +50 mV (Figures 4e, f, h and Supplementary Figures 21–23). There were also some channels with conductances between 0.9 and 1.2 nS. This last population may be nanotube seed pores or nanotubes without attached channel caps (Figure 4f, h). Like the DNA seed channels and the nanotube seed channels, the seed channels with attached channel caps exhibited an ohmic conductance pattern (Figure 4g). To verify that the binding of the channel caps was responsible for the reduction in observed conductance of the seed pores, we removed the adapter strands that mediate the binding of the channel caps to the seeds to make “control” channel caps. These control channels could not bind to the seed pores, but were added to the membranes along with the seed pores (see Materials and Methods, Supplementary Notes 5 and 11). Fluorescence micrographs showed little or no interaction between seeds and control channel caps after they were incubated together (Supplementary Figures 24a and d). The conductances observed across lipid bilayers using a mixture of nanotube seed channels and the control channel caps were the same as those of the seed channels alone, *i.e*., around 1–1.2 nS (Supplementary Figures 24b and c). Thus, the ability of the seed channel and channel caps to bind is required for the drop in the conductance of the channel observed. These experiments further imply that the pores remain bound to the channel caps in the presence of transmembrane potential. Interestingly, no gating of the seed pore channel bound to channel caps was observed. The lack of gating and the constant conductance of the channels we observed suggests that nanotube seed channels and channel caps bind irreversibly and do not change conformation over the timescales while inserted into the channel.

The conductances of 0.4 and 0.6 nS produced by seed-channel cap complexes are comparable to that of the 6-helix DNA bundle constructed by Mendoza *et al*. Their DNA helical nanopore produces a conductance of about 0.3 nS, whose structure has a lock strand attached to the barrel. Upon addition of the key strands, the DNA channel produces a higher conductance of 0.5 nS, presumably due to the widening of the channel aperture.^58^ Comparable conductance changes are also observed in a six-helix bundle nanopore^29^ when a lock strand binds to the channel entrance, causing a reduction in ion-flow. Here we show that the binding of a channel cap complex to the seed channel, which reduced the size of the channel’s diameter, can likewise reduce channel conductance.^38^ The channel cap’s reduction of the conductance by only about a factor of two suggests that there is ionic flow through the walls of the DNA channel.^56,59^

The mean and standard deviation of the conductances of open DNA nanotube channels, which have an average length of 2.43 ± 0.10 μm (95% CI, N = 185, Supplementary Figure 11), were similar to the mean and standard deviation of the conductances of the nanotube seeds, which are monodisperse with a length of 64.70 ± 2.12 nm, entirely inconsistent with the predictions of the theoretical model that assumes ion flow from channel end to channel end: this model predicts that conductance decreases significantly with channel length. The similarity between the conductances of the nanotube seed and the nanotube channel therefore also suggests that a significant amount of ion flow occurs through seed, and presumably also nanotube, channel walls. We next used the channel cap to characterize the degree of ion flow through the walls of the nanotubes, and therefore the extent to which ions can travel from one end of the nanotube through the membrane to the other end. The same channel cap designed to bind to a nanotube seed can also bind to the end of a DNA nanotube, as the free end of a DNA nanotube presents the same sticky ends as their nanotube seed templates (Figure 1). Nanotubes with channel caps were assembled by first assembling seeded nanotubes using the same method as in previous experiments and then incubating the nanotubes with the channel caps at 32°C for 3 h (see Materials and Methods, Figure 5a and Supplementary Figure 10). The nanotubes with the channel caps were further incubated with the DNA-cholesterol conjugate for 1 h under ambient conditions and were then centrifuged at 3000 g for 30 min for 2–3 times using the 1X TAEM buffer supplemented with 12.5 mM magnesium chloride. After filter purification, 82 ± 3% of the structures in fluorescence micrographs were seeded nanotubes with attached channel caps, 9 ± 4% of the structures observed were DNA seed structures with attached channel caps, and the remaining 3 ± 2% and 4 ± 3% of structures observed were seeds and seeded tubes without caps (Figure 5b and Supplementary Figure 25a). Thus >90% of structures had channel caps. The average length of the capped nanotubes before purification was 1.87 ±0.10 μm (95% CI, N = 108) and 1.57 ± 0.20 (95% CI, N = 105) after purification, confirming that purification did not lead to a large change in the lengths of the DNA nanotube structures (Supplementary Figure 25b). The resulting nanotube channels with caps produced changes in conductance across DPhPC bilayers, suggesting that they could insert into lipid bilayers (Figure 5c–e). The structures showed a much broader range of conductances than the other structures (Figure 5f and Supplementary Figures 26 and 27). We observed three different forms of conductance, namely, 396 ± 45 pS, 697 ± 70 pS and 1158 ± 172 pS, although the small number of samples make it difficult to be confident in the precise location and number of these peaks. The existence of multiple peaks is consistent with the fact that the conductance of a combination of structures (seed pores and nanotube channels with and without caps) could insert into the membrane. The peak (1158 ± 172 pS) with the same conductance as those of open nanotubes accounts for at least 30% of the measurements, even though <10% of the structures being characterized are uncapped nanotube channels. This suggests that a fraction of capped nanotube channels had similar conductances to the open ones. Perhaps these nanotubes have rough end facets, which would create gaps in their binding interface with channel caps. These holes would be the size of nanotube monomers (about 13 × 4 nm) so that the channel cap does not effectively reduce the size of a channel’s opening. The other peaks are similar in magnitude to the conductances of nanotube seed-channel cap complexes. These results suggest that ion transport in at least some cases proceed from one end of a microns-long nanotube to the other across the membrane.

**Figure 5.**
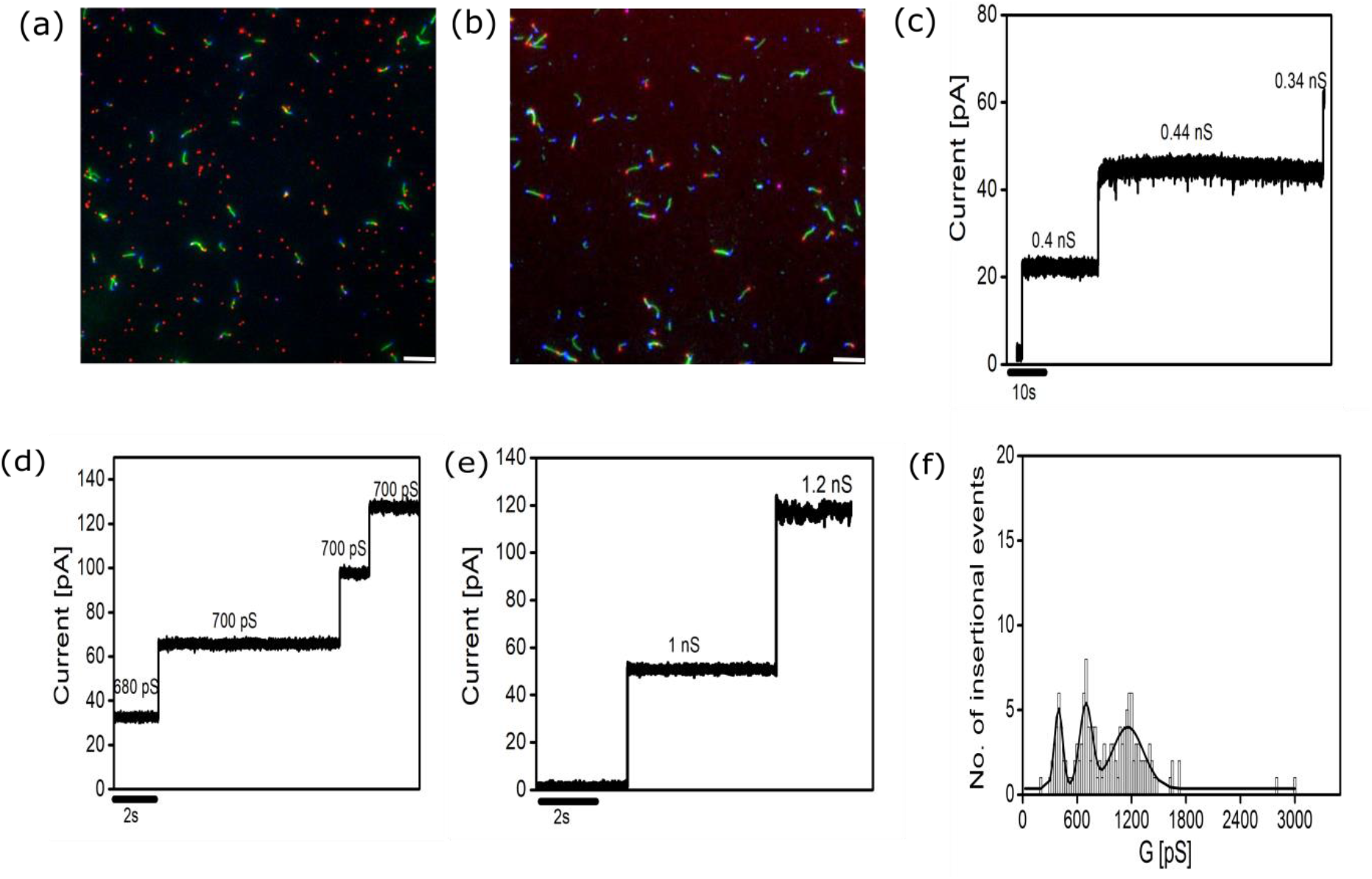
Conductivity across capped DNA nanotube membrane channels. (a) Fluorescence micrographs of capped nanotubes (a) before cholesterol-DNA conjugates were added and (b) after cholesterol-DNA conjugates were added and the structures were purified by 2 rounds of centrifugation at 3000 g for 30 min. Seeds are labeled with ATTO488 (blue), channel caps are labeled with ATTO647N (red) and nanotubes are labeled with Cy3 (green). Scale bars: 5 μm. (c–e) Electrical signatures showing different conductance patterns exhibited by capped seeded DNA nanotube channel in 1 M KCl, 10 mM MgCl_2_, buffered with 10 mM MES, at pH 6.0 at an applied transmembrane potential of +50 mV. (f) Histogram of the conductance steps of capped DNA nanotube channels across DPhPC bilayers formed by the DIB technique at an applied transmembrane potential of +50 mV (N = 144). A Gaussian fit to the conductances is shown as an overlay of three different forms of conductances in the range of 396 ± 45 pS, 697 ± 70 pS and 1158 ± 172 pS.

## CONCLUSIONS

DNA nanostructures, which can be designed to have specific shapes by design, in combination with hydrophobic molecules, can be used to form a range of transmembrane pores by design.^22–31^ In this paper, we explore the use of DNA nanostructures to form nanofluidic channels that might also transport ions or other species along pathways microns in length by characterizing how ions flow through these structures when an electrical potential is applied.

The channels that we have constructed are ohmic in character and have conductances similar to analogous channels published in previous reports.^22–24,27,29,30^ The reduced conductance of both nanotube seed and nanotube channels when channel caps are attached as compared to their uncapped counterparts suggests that the ions transported through these channels moved from one cylindrical end of the channel to the other and through channel walls. These results support the idea that DNA nanostructures might be used to direct ion flow across distances of many microns, and also suggest ways that mechanisms that might reduce transport through walls, such as coating the nanotube’s surface, might be quantitatively assessed. The ability to readily assemble DNA nanotubes under physiological conditions^40^ or to target or orient the terminals at the sites of particular biomolecular labels^39^ means that such structures might be directed to form interconnects or channels at specific locations on cells or synthetic devices. The ability of nanotubes to direct the flow of ions, measured via conductivity across the membrane, suggests a means for the assembly of regenerative or dynamic bio-electronic devices with well-defined circuit topologies. Coating such structures with lipids^60,61^, proteins^62–64^, polycations^65^/peptoids^66^, block co-polymers^67^ or silica shells^68^ might allow ion-transport through channels with no ion loss across channel walls. Ion transport within insulated biomolecular channels is a key means of charge transport in living systems. The ability to recapitulate this means of transport in engineered systems could be a useful means of building synthetic, self-assembled circuits for rapidly transporting information or chemical signals via charge transport.

## Supporting Information

Additional electrical recordings for nanotube seeds, nanotubes, nanotube seed-channel cap complex, capped nanotubes, fluorescence micrographs of the seeded nanotubes, nanotube seed-channel complex, capped nanotubes, additional transmission electron micrographs of the the DNA origami nanostructures, detailed experimental protocol for the construction of nanotube seeds, nanotubes, nanotube seed-channel cap complex, capped nanotubes and complete sequences of the nanotube seed, nanotubes and channel cap.

## ACKNOWLEDGMENTS

The authors thank Andy Sarles and Joseph Najem for their help in droplet-interface bilayer measurements. This work was supported by DARPA YFA D16AP00147 to Rebecca Schulman.

## METHODS

### DNA strands

The sequences for nanotube seeds and nanotubes are listed in the Supplementary Information (Supplementary Notes 1 and 2 and Supplementary Figures 1–3). All the DNA strands and the DNA-cholesterol conjugate used in this work were purchased from Integrated DNA technologies (IDT, Iowa, USA), except the M13mp18 scaffold strand which was purchased from Bayou Biolabs (Los Angeles, USA). The strands that comprise the DNA nanotube monomers and the DNA-cholesterol conjugate were HPLC purified; the adapter strands on the nanotube seeds and channel caps that act as binding sites for monomers or nanotubes were PAGE purified. The staple strands for the nanotube seed and channel cap were not purified. The concentrations of the DNA tiles, adapters and the cholesterol strands were determined by measuring the absorbance at 260 nm using a spectrophotometer (Eppendorf Model 6131, Hamburg, Germany) and the Beer-Lambert law using the extinction coefficients provided by IDT.

### Materials

Potassium chloride (Alfa Aesar, Massachusetts, USA. Product No: 10220196), 4-morpholineethanesulfonic acid (MES) monohydrate (Alfa Aesar, Massachusetts, USA. Product No: 10191845), calcium chloride (Sigma Aldrcih, USA. Product No: SLBW7125), magnesium chloride (Sigma Aldrich, USA. Product No: SLB3227V), hexadecane (Sigma Aldrich, USA. Product No: SHBK6702), TAE buffer (Ultrapure™ DNA typing grade^®^, Invitrogen, USA), sodium hypochlorite (Sigma Aldrich, USA), agarose tablets (EZ Pack Agarose, Benchmark Scientific, Inc, New Jersey, USA. Product No: A2501), uranyl acetate (Electron Microscopy Sciences, Philadelphia, USA. Product No: 22400), bovine serum albumin (Sigma Aldrich, Product No: SLBT2162) and Amicon^®^ Ultra centrifugal filters 100 kDa (100 K, Merck Millipore Ltd., Darmstadt, Germany).

#### Sample preparation

##### DNA nanotube seeds and nanotube seed channels

Seeds were annealed as described in Mohammed *et al*, 2013.^38^ Seeds used to grow nanotubes and seeds were annealed in 40 mM Tris-acetate, 1 mM EDTA supplemented with 12.5 mM magnesium acetate, pH 8.0 (1X TAEM buffer). To make DNA seed channels, the DNA-cholesterol conjugate strand (3’ nanotube seed_Chol.DNA, see Supplementary Note 1 for sequences) dissolved in MilliQ water was heated to 50°C in an Eppendorf tube for 15 min to dissolve agglomerates. The DNA conjugate strand was then added to a final concentration of 10 μM to 50 μL of unpurified DNA seed origami solution. The solution was then incubated for 1 h under ambient conditions. For electrical measurements, after incubation of the nanotube seeds with the DNA-cholesterol conjugates, the DNA seed structures were then purified from excess staples and unhybridized cholesterol conjugates via spin filtration using 100 kDa MWCO filters (Amicon^®^ ultra centrifugal filters – 100 K) in 350 μL 1X TAEM buffer. The filtration process was repeated at 3000 g for 30 min at least 3–5 times (Supplementary Notes 4 and 6).

For control experiments where no DNA-cholesterol conjugate was added to the DNA nanotube seeds, purified seeds were directly mixed with lipid vesicles prior to electrophysiology experiments. The conductance of 10 μM of the DNA-cholesterol conjugates alone was also measured as a control, by directly mixing the cholesterol strands with the lipid vesicles and then performing the electrical measurements.

##### DNA nanotubes

The strands for a) the nanotube seeds and b) 40 nM of the nanotube tiles were annealed in 1X TAEM buffer in two separate tubes. The tubes were initially heated at 90°C for 5 min to melt all DNA, cooled from 90 to 45°C at 1°C/min and then incubated at 45°C for 1 h to allow the formation of seeds and nanotube tiles. The samples were then slowly cooled from 45 to 32°C at 0.1°C/min to a final temperature of 32°C. The annealed seed structures were then spin filtered using 100 kDa MWCO filters (Amicon^®^ ultra centrifugal filters – 100 K). The spin filtration process was repeated at 3000 g for 4 min at least 4 times (no addition of 3’ nanotube seed_Chol.DNA). The purified seeds were then added to a final concentration of 18 pM to a mixture of 40 nM of nanotube tiles mix at 32°C. The reaction mixture was then incubated at 32°C for 18–24 h or longer to allow seeded nanotubes to grow.^38^ To make DNA nanotube channels, DNA-cholesterol conjugate solution was heated as described for nanotube seed channel preparation, then added to a final concentration of 10 μM to a solution of 18 pM of seeded DNA nanotubes. DNA nanotube channels were filter purified as described for nanotube seed channels. Before use in lipid bilayer experiments, the nanotubes were imaged using fluorescence microscopy to confirm the percentage of nanotubes seeds with tubes and the approximate tube length. For control experiments with the DNA nanotubes alone (no seeds attached), 250 nM of each strand of the DNA tiles were annealed following the same protocol as that of the seeded tubes and the reaction mixture was kept at 32°C for 24 h. The DNA tubes were further filter purified at 3000 g for 4 min four times and directly mixed with DPhPC vesicles prior to electrophysiology experiments (Supplementary Note 8).

##### Channel caps and nanotube seed channels with attached channel caps

Channel caps were annealed and filter purified following the methods for nanotube seed annealing and purification (Supplementary Note 9). Nanotube seed channels with attached channel caps were prepared by mixing nanotube seeds and channel caps at a stoichiometric ratio of 1:3. The mixture was then incubated for approximately 3 h at 32°C. To the incubated sample, pre-heated DNA-cholesterol conjugates were added to a final concentration of 10 μM and further incubated for 1 h under ambient conditions. The resulting structures were filter purified for 30 min at 3000 g for 2–3 times.

##### Shuttered DNA nanotube channels

Seeded DNA nanotubes were prepared and filter purified as described above. Seeded nanotubes were mixed with channel caps at the same stoichiometric ratio of nanotube–seeds:channel–caps as for seed channels with channel caps and were incubated with DNA-cholesterol conjugates then filter purified in the same manner as nanotube seed channels with channel caps (Supplementary Note 10).

##### Transmission electron microscopy

Formvar-coated carbon grids (Electron Microscopy Sciences, Philadelphia, USA) were used for imaging DNA nanostructures. Prior to imaging the DNA nanostructures, the grids were glow discharged for 40 s. To image nanotube seeds or channel caps, 5–10 μL of a solution containing the seeds (1.2 nM) or channel caps (0.4 nM) was added directly onto the grid, left for 10 min and then washed with MilliQ water. To image the nanotube seeds bound to channel caps, the nanotube seeds and the channel caps were prepared and then pre-incubated together before at 32°C for about 3 h prior to imaging under TEM. The grids were then stained with 50 μL aqueous uranyl acetate solution (2% v/v) (Electron Microscopy Sciences, Philadelphia, USA. Product No: 22400) for 5 min. The excess staining solution was carefully removed using filter paper. The grids were then dried in a vacuum chamber for 3 h prior imaging using a field emission Tecnai-12 TEM with an operating voltage at 100 keV.

##### Preparation of small unilamellar vesicles (SUV’s)

1,2-diphytanoyl-*sn*-glycero-3-phosphocholine (DPhPC, Avanti Polar Lipids, USA) was used to form small unilamellar vesicles (SUV’s). Briefly, the lipids were dissolved in chloroform in a glass vial at a concentration of 5 mg/mL. The glass vial was kept under nitrogen flow for 20 min to completely evaporate the chloroform. The lipid film that formed was then kept under vacuum in a vacuum chamber for 24 h. The lipid film was then hydrated by resuspending the sample mixture in 1 mL of 150 mM potassium chloride in water. SUV’s were then formed by sonicating the solution in a bath sonicator (Bransonic Ultrasonic Cleaner (1510R-DTH)) for 2 h until the solution turned completely transparent. Prior to imaging, the nanotube seed channels or the nanotube channels with the SUV’s, the SUV’s were diluted 100 times with 150 mM of potassium chloride buffer. The diluted SUV’s were then mixed either with the DNA seed pore or the seeded nanotube pore solution prepared as described above at a volumetric ratio of 1:3.

##### Electron microscopy imaging of the nanotube seeds and DNA nanotubes with SUV’s

Formvar coated carbon grids (Electron Microscopy Sciences, Philadelphia, USA) were used for imaging the interaction of DNA nanostructures with SUV’s. Prior to imaging, the grids were glow discharged for 40 s. To characterize the interaction of nanotube seeds or nanotubes with SUV’s via TEM, 10 μL of a solution containing the DNA nanostructure samples with SUV’s in the ratio 1:3 was added directly onto a grid and left for 10 min, then washed with MilliQ water. The grids were then stained with 50 μL aqueous uranyl acetate solution (2% v/v). The excess staining solution was carefully removed using filter paper. The grids were then dried in a vacuum chamber for 3 h prior imaging using a FEI Tecnai-12 with an operating voltage at 100 keV.

##### Preparation of phospholipid vesicles for electrical measurements

DPhPC was dissolved in chloroform at a concentration of 5 mg/mL in a glass vial. The glass vial was kept under vacuum for 16 h to allow the chloroform to evaporate. A stream of nitrogen was then flowed through the sample to remove the residual solvent. The lipid film that formed was then resuspended in an aqueous solution containing 1 M potassium chloride, 10 mM MgCl_2_ and 10 mM MES monohydrate at pH 6.0. The solution was freeze-thawed at least 6 times under liquid nitrogen and then extruded 21 times using an Avanti pressure extruder with Whatmanfilter paper of size 0.1 μm (Avanti Polar Lipids Alabama, USA. Product No: SKU 640005-1Ea). The resulting transparent solution was then used for lipid bilayer experiments.

##### Single-channel electrical recordings

All electrical measurements were performed using the droplet interface bilayer technique.^48–50^ The aqueous phase of the droplets consisted of phospholipid vesicles prepared using 1 M KCl, 10 mM MES, 10 mM MgCl_2_, pH 6.0; hexadecane acted as the oil phase. Briefly, one end of the silver wires (diameter – 125 μm, WPI, USA) was melted over a fire flame to create a ball end. The ball end of the silver wire was then bleached for 30 min using 20% sodium hypochlorite solution. The ball end of the silver wire was then dipped into a solution of 1% agarose dissolved in water. To avoid dehydration, the agarose-coated silver wire was then placed immediately in the chamber containing hexadecane. Approximately 0.2 μL droplet of the lipid vesicle solution was added to agarose-coated region of the silver electrodes. The two lipid droplets residing in the hexadecane oil phase led to the formation of a stable monolayer. After 15–20 min, the electrodes were then brought in close proximity to allow the lipid droplets to come into contact with each other. Capacitance was measured by monitoring the square wave output current upon applying a triangular wave input voltage (10 mV/10 Hz frequency) across the bilayer. DNA nanostructures were then added to one of the droplets and a transmembrane potential was applied to induce DNA nanopore insertion. Electrical recording was performed using an Axopatch 200B and the Digidata 1440A (Molecular Devices, USA). The signals were recorded at 1 kHz and sampled at 50 Hz with an 8-pole Bessel filter and directly saved into computer memory. The conductance data was analyzed using Clampfit 10.7 software and the graphs were plotted using the Origin Pro 8.0.

## Table of Contents graphic

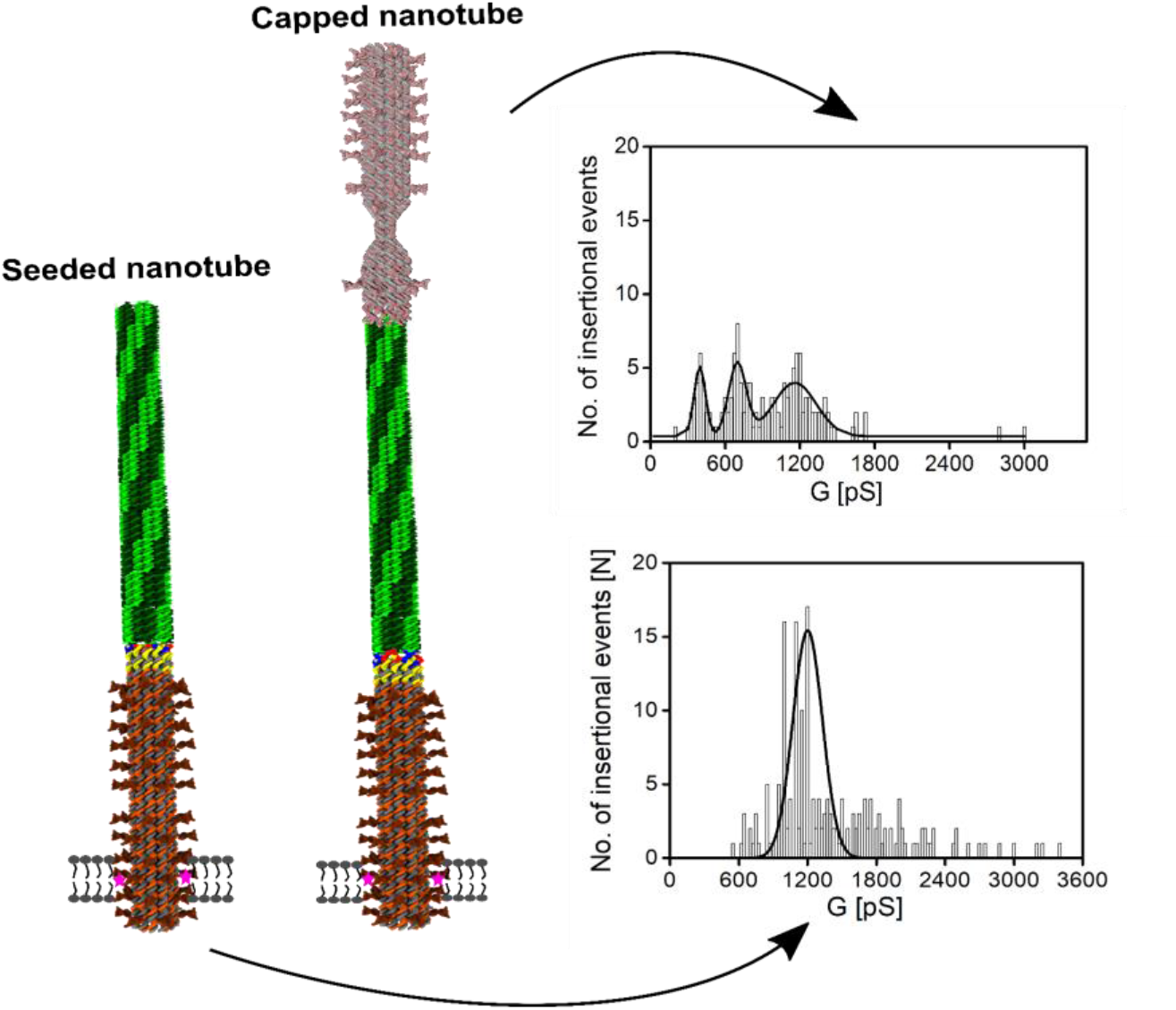

## REFERENCES

(1) Evans, W. H.; Martin, P. E. Gap junctions: structure and function. Mol. Membr. Biol. 2002, 19, 121–136.

(2) Doyle, D. A.; Morais Cabral, J.; Pfuetzner, R. A.; Kuo, A.; Gulbis, J. M.; Cohen, S. L.; Chait, B. T.; MacKinnon, R. The structure of the potassium channel: molecular basis of K+ conduction and selectivity. Science 1998, 280, 69–77.

(3) Ferrara, L. G. M.; Wallat, G. D.; Moynié, L.; Dhanasekar, N. N.; Aliouane, S.; Acosta-Gutiérrez, S.; Pagès, J. M.; Bolla, J. M.; Winterhalter, M.; Ceccarelli, M.; Naismith, J. H. MOMP from Campylobacter jejuni is a trimer of 18-stranded β-barrel monomers with a Ca^2+^ ion bound at the constriction zone. J. Mol. Biol. 2016, 428, 4528–4543.

(4) Mantri, S.; Sapra, K. T.; Cheley, S.; Sharp, T. H.; Bayley, H. An engineered dimeric protein pore that spans adjacent lipid bilayers. Nat. Commun. 2013, 4.

(5) Sowinski, S.; Jolly, C.; Berninghausen, O.; Purbhoo, M. A.; Chauveau, A.; Köhler, K.; Oddos, S.; Eissmann, P.; Brodsky, F. M.; Hopkins, C.; Önfelt, B.; Sattentau, Q.; Davis, D. M. Membrane nanotubes physically connect T cells over long distances presenting a novel route for HIV-1 transmission. Nat. Cell. Biol. 2008, 10, 211–219.

(6) La Boissiere, S.; Izeta, A.; Malcomber, S.; O’Hare, P. Compartmentalization of VP16 in cells infected with recombinant herpes simplex virus expressing VP16-green fluorescent protein fusion proteins. J. Virol. 2004, 78, 8002–8014.

(7) Gerdes, H.-H. Prions tunnel between cells. Nat. Cell. Biol. 2009, 11, 235.

(8) Kasianowicz, J.J.; Brandin, E.; Branton, D.; Deamer, D.W. Characterization of individual polynucleotide molecules using a membrane channel. Proc. Natl. Acad. Sci. USA 1996, 93, 13770–13773.

(9) Deamer, D.; Akeson, M.; Branton, D. Three decades of nanopore sequencing. Nat. Biotechnol. 2016, 34, 518–524.

(10) Wloka, C.; Meervelt, V. V.; van Gelder, D.; Danda, N.; Jager, N.; Williams, C. P.; Maglia, G. Label-free and real-time detection of protein ubiquitination with a biological nanopore. ACS Nano 2017, 11, 4387–4394.

(11) Soskine, M.; Biesemans, A.; Moeyaert, B.; Cheley, S.; Bayley, H.; Maglia, G. An engineered ClyA nanopore detects folded target proteins by selective external association and pore entry. Nano Lett. 2012, 12, 4895–4900.

(12) Galenkamp, N. S.; Biesemans, A.; Maglia, G. Directional conformer exchange in dihydrofolate reductase revealed by single-molecule nanopore recordings. Nat. Chem. 2020, 12, 481–488.

(13) Harrington, L.; Alexander, L. T.; Knapp,S.; Bayley, H. Single-molecule protein phosphorylation and dephosphorylation by nanopore enzymology. ACS Nano 2019, 13, 633–641.

(14) Fahie, M.; Chisholm, C.; Chen, M. Resolved single-molecule detection of individual species within a mixture of anti-biotin antibodies using an engineered monomeric nanopore. ACS Nano 2015, 9, 1089–1098.

(15) Wang, Y.; Tian, K.; Shi, R.; Gu, A.; Pennella, M.; Alberts, L.; Gates, K. S.; Li, G.; Fan, H.; Wang, M. X.; Gu, L-Q. Nanolock-nanopore facilitated digital diagnostics of cancer driver mutation in tumor tissue. ACS Sensors 2017, 2, 975–981.

(16) Liu, P.; Kawano, R. Recognition of single-point mutation using a biological nanopore. Small Methods 2020.

(17) Booth, M. J.; Cazimoglu, I.; Bayley, H. Controlled deprotection and release of a small molecule from a compartmented synthetic tissue module. Commun. Chem. 2019, 2.

(18) Downs, F. G.; Lunn, D. J.; Booth, M. J.; Sauer, J. B.; Ramsay, W. J.; Klemperer, R. G.; Hawker, C. J.; Bayley, H. Multi-responsive hydrogel structures from patterned droplet networks. Nat. Chem. 2020, 12, 363–371.

(19) Gerdes, H.-H.; Rustom, A.; Wang, X. Tunneling nanotubes, an emerging intercellular communication route in development. Mech. Dev. 2013, 130, 381–387.

(20) Abounit, S.; Zurzolo, C. Wiring through tunneling nanotubes – from electrical signals to organelle transfer. J. Cell. Sci. 2012, 125, 1089–1098.

(21) Schild, V. R.; Booth, M. J.; Box, S. J.; Olof, S. N.; Mahendran, K. R.; Bayley, H. Light-patterned current generation in a droplet bilayer array. Sci. Rep. 2017, 7, 46585.

(22) Langecker, M; Arnaut, V; Martin, T. G.; List, J; Renner, S; Mayer, M; Dietz, H; Simmel, F. C. Synthetic lipid membrane channels formed by designed DNA nanostructures. Science 2012, 338, 932–936.

(23) Burns, J. R.; Stulz, E. Howarka S. Self-assembled DNA nanopores that span lipid bilayers. Nano Lett. 2013, 13, 2351–2356.

(24) Burns, J. R.; Göpfrich, K.; Wood, J. W.; Thacker, V. V.; Stulz, E.; Keyser, U. F.; Howarka, S. Lipid-bilayer-spanning DNA nanopores with a bifunctional porphyrin anchor. Angew. Chem. Int. Ed. 2013, 52, 12069–12072.

(25) Krishnan, S.; Ziegler, D.; Arnaut, V.; Martin, T. G.; Kapsner, K.; Henneberg, K.; Bausch, A. R.; Dietz, H.; Simmel, F. C. Molecular transport through large-diameter DNA nanopores. Nat. Commun. 2016, 7, 12787.

(26) Göpfrich, K; Li, C. Y.; Ricci, M.; Bhamidimarri, S. P.; Yoo, J.; Gyenes, B.; Ohmann, A.; Winterhalter, M.; Aksimentiev, A.; Keyser, U. F. Large-conductance transmembrane porin made from DNA origami. ACS Nano 2016, 10, 8207–8214.

(27) Diederichs, T.; Pugh, G.; Dorey, A.; Xing, Y.; Burns, J. R.; Hung Nguyen, Q.; Tornow, M.; Tampé, R.; Howorka, S. Synthetic protein-conductive membrane nanopores built with DNA. Nat. Commun. 2019, 10, 5018.

(28) Lv, C.; Gu, X.; Li, H.-W.; Zhao, Y.; Yang, D.; Yu, W.; Han, D.; Li, J.; Tan, W. Molecular transport through a biomimetic DNA channel on live cell membranes. ACS Nano 2020, 14, 14616–14626.

(29) Burns, J. R.; Seifert, A.; Fertig, N.; Howorka, S. A biomimetic DNA-based channel for the ligand-controlled transport of charged molecular cargo across a biological membrane. Nat. Nanotechnol. 2016, 11, 152–156.

(30) Seifert, A.; Göpfrich, K.; Burns, J. R.; Fertig, N.; Keyser, U. F.; Howorka, S. Bilayer-spanning DNA nanopores with voltage-switching between open and closed state. ACS Nano 2015, 9, 1117–1126.

(31) Göpfrich, K.; Zettl, T.; Meijering, A. E. C.; Hernańdez-Ainsa, S.; Kocabey, S.; Liedl, T.; Keyser, U. F. DNA-tile structures induce ionic currents through lipid membranes. Nano Lett. 2015, 15, 3134–3138.

(32) Burns, J. R.; Al-Juffali, N.; Janes, S. M.; Howorka, S. Membrane-spanning DNA nanopores with cytotoxic effect. Angew. Chem. Int. Ed. Engl. 2014, 53, 12466–12470.

(33) Zhang, Q.; Jiang, Q.; Li, N.; Dai, L.; Liu, Q.; Song, L.; Wang, J.; Li,Y.;Tian, J.; Ding, B.; Du, Y. DNA origami as an *in vivo* drug delivery vehicle for cancer therapy. ACS Nano 2014, 8, 6633–6643.

(34) Jiang, Q.; Liu, S.; Liu, J.; Wang, Z.-G.; Din, B. Rationally designed DNA-origami nanomaterials for drug delivery in vivo. Adv. Mater. 2018, 31, e1804785.

(35) Douglas, S. M.; Bachelet, I.; Church, G. M. A logic-gated nanorobot for targeted transport of molecular payloads. Science 2012, 335, 831–834.

(36) Zhao, Y-X.; Shaw, A.; Zeng, X.; Benson, E.; Nyström, A. M.; Högberg, B. DNA origami delivery system for cancer therapy with tunable release properties. ACS Nano 2012, 6, 8684–8691.

(37) Wang, D.; Zhang, Y.; Wang, M.; Dong, Y.; Zhou, C.; Isbell, M. A.; Yang, Z.; Liu, H.; Liu, D. A switchable DNA origami nanochannel for regulating molecular transport at the nanometer scale. Nanoscale 2016, 8, 3944–3948.

(38) Mohammed, A. M.; Schulman, R. Directing self-assembly of DNA nanotubes using programmable seeds. Nano Lett. 2013, 13, 4006–4013.

(39) Mohammed, A. M.; Šulc, P.; Zenk, J.; Schulman, R. Self-assembling DNA nanotubes to connect molecular landmarks. Nat. Nanotechnol. 2017, 12, 312–316.

(40) Li, Y.; Schulman, R. DNA nanostructures that self-heal in serum. Nano Lett. 2019, 19, 3751–3760.

(41) Agrawal, D. K.; Jiang, R.; Reinhart, S.; Mohammed, A. M.; Jorgenson, T. D.; Schulman, R. Terminating DNA tile assembly with nanostructured caps. ACS Nano 2017, 11, 9770–9779.

(42) Dietz, H.; Douglas, S. M.; Shih, W. M. Folding DNA into twisted and curved nanoscale shapes. Science 2009, 325, 725–730.

(43) Douglas, S.M.; Dietz, H.; Liedl, T.; Hogberg, B.; Graf, F.; Shih, W.M. Self-assembly of DNA into nanoscale three-dimensional shapes. Nature 2009, 459, 414–418.

(44) Ren, R.; Zhang, Y.; Nadappuram, B. P.; Akpinar, B.; Klenerman, D.; Ivanov, A. P.; Edel, J. B.; Korchev, Y. Nanopore extended field-effect transistor for selective single-molecule biosensing. Nat. Commun. 2017, 8, 586.

(45) Al Nahas, K.; Cama, J.; Schaich, M.; Hammond, K.; Deshpande, S.; Dekker, C.; Ryadnov, M. G.; Keyser, U. F. A microfluidic platform for the characterisation of membrane active antimicrobials. Lab. Chip. 2019, 19, 837–844.

(46) Schaich, M.; Cama, J.; Al Nahas, K.; Sobota, D.; Sleath, H.; Jahnke, K.; Deshpande, S.; Dekker, C.; Keyser, U. F. An integrated microfluidic platform for quantifying drug permeation across biomimetic vesicle membranes. Mol. Pharm. 2019, 16, 2494–2501.

(47) Rothemund, P. W. K. Folding DNA to create nanoscale shapes and patterns. Nature 2006, 440, 297–302.

(48) Bayley, H.; Cronin, B.; Heron, A.; Holden, M. A.; Ruhma, W.; Syeda, R.; Thompson, J.; Wallace, M. Droplet interface bilayers. Mol. Biosyst. 2008, 4, 1191–1208.

(49) Leptihn, S.; Castell, O. K.; Cronin, B.; Lee, E.-H.; Gross, L. C. M.; Marshall, D. P.; Thompson, J. R.; Holden, M.; Wallace, M. I. Constructing droplet interface bilayers from the contact of aqueous droplets in oil. Nat. Protocol. 2013, 6, 1048–1057.

(50) Najem, J. S.; Dunlap, M. D.; Rowe, I. D.; Freeman, E. C.; Grant, J. W.; Sukharev, S.; Leo, D. J. Activation of bacterial channel MscL in mechanically stimulated droplet interface bilayers. Sci. Rep. 2015, 5, 13726.

(51) Hille, B. Ion channels of excitable membranes, 3^rd^ ed.; Sinauer Associates: Sunderand MA, USA, 2001.

(52) Ho, C.; Qiao, R.; Heng, J. B.; Chatterjee, A.;Timp, R. J. Electrolytic transport through a synthetic nanometer-diameter pore. Proc. Natl. Acad. Sci. USA 2005, 102, 10455–10450.

(53) Geng, J.; Kim, K.; Zhang, J.; Escalada, A.; Tunuguntla, R.; Comolli, L. R.; Allen, F. I.; Shnyrova, A. V.; Cho, K. R.; Munoz, D.; Wang, Y. M.; Grigoropoulos, C. P.; Ajo-Franklin, C. M.; Frolov, V. A.; Noy, A. Stochastic transport through carbon nanotubes in lipid bilayers and live cell membranes. Nature 2014, 514, 612–615.

(54) Maingi, V.; Lelimousin, M.; Howorka, S; Sansom, M. S. Gating-like motions and wall porosity in a DNA nanopore scaffold revealed by molecular simulations. ACS Nano 2015, 9, 11209–11217.

(55) Maingi, V.; Burns, J. R.; Uusitalo, J. J.; Howorka, S.; Marrink, S. J.; Sansom, M. S. P. Stability and dynamics of membrane-spanning DNA nanopores. Nat. Commun. 2016, 8, 14784.

(56) Yoo, J.; Aksimentiev, A. Molecular dynamics of membrane-spanning DNA channels: conductance mechanism, electro-osmotic transport, and mechanical gating. J. Phys. Chem. Lett. 2015, 6, 4680–4687.

(57) Yoo. J.; Aksimentiev, A. In situ structure and dynamics of DNA origami determined through molecular dynamics simulations. Proc. Natl. Acad. USA 2013, 110, 20099–20104.

(58) Mendoza, O.; Calmet, P.; Alves, I.; Lecomte, S.; Raoux, M.; Cullin, C.; Elezgaray, J. A tensegrity driven DNA nanopore. Nanoscale 2017, 9, 9762–9769.

(59) Plesa, C.; Ananth, A.N.; Linko, V.; Gülcher, C.; Katan, A.J.; Dietz, H.; Dekker, C. Ionic permeability and mechanical properties of DNA origami nanoplates on solid-state nanopores. ACS Nano 2014, 8, 35–45

(60) Perrault, S. D.; Shih, W. M. Virus-inspired membrane encapsulation of DNA nanostructures to achieve *in vivo* stability. ACS Nano 2014, 8, 5132–5140.

(61) Perrault, S. D.; Shih, W. M. Lipid membrane encapsulation of a 3D DNA nano octahedron. Meth. Mol. Biol. 2017, 1500, 165–184.

(62) Auvinen, H.; Zhang, H.; Nonappa, Kopilow, A.; Niemelä, E. H.; Nummelin, S.; Correia, A.; Santos, H. A.; Linko, V.; Kostiainen, M. A. Protein coating of DNA nanostructures for enhancedstability and immunocompatibility. Adv. Healthcare. Mater. 2017, 6, 1700692–1700698.

(63) Ponnuswamy, N.; Bastings, M. M. C.; Nathwani, B.; Ryu, J. H.; Chou, L. Y. T.;Vinther, M.; Li, W. A.; Anastassacos, F. M.; Mooney, D. J.; Shih, W. M. Oligolysine-based coating protects DNA nanostructures from low-salt denaturation and nuclease degradation. Nat. Commun. 2017, 8, 15654.

(64) Anastassacos, F. M.; Zhao, Z.; Zeng, Y.; Shih, W. M. Glutaraldehyde cross-linking of oligolysines coating DNA origami greatly reduces susceptibility to nuclease degradation. J. Am. Chem. Soc. 2020, 142, 3311–3315.

(65) Ahmadi, Y.; De Llano, E.; Barišić, I. (Poly)cation-induced protection of conventionaland wireframe DNA origami nanostructures. Nanoscale 2018, 10, 7494–7504.

(66) Wanga, S.-H.; Gray, M. A.; Xuan, S.; Lin, Y.; Byrnes, J.; Nguyen, A. I.; Todorova, N.; Stevens, M. M.; Bertozzi, C. R.;Zuckermann, R. N.; Gang, O. DNA origami protection and molecular interfacing through engineered sequence-defined peptoids. Proc. Natl. Acad. Sci. USA 2020, 117, 6339–6348.

(67) Agarwal, N. P.; Matthies, M.; Gür, F. N.; Osada, K.; Schmidt, T. L. Block copolymer micellization as a protection strategy for DNAorigami. Angew. Chem. Int. Ed. Engl. 2017, 56, 5460–5464.

(68) Liu, X.G.; Zhang, F.; Jing, X.X.; Pan, M.C.; Liu, P.; Li, W.; Zhu, B.W.; Li, J.; Chen, H.; Wang, L.H.; Lin, J.P.; Liu, Y.; Zhao, D.Y.; Yan, H.; Fan, C.H. Complex silica composite nanomaterials templated with DNA origami. Nature 2018, 559, 593–598.

