## Supplemental Figure 1 for "The ion permeability of DNA nanotube channels"

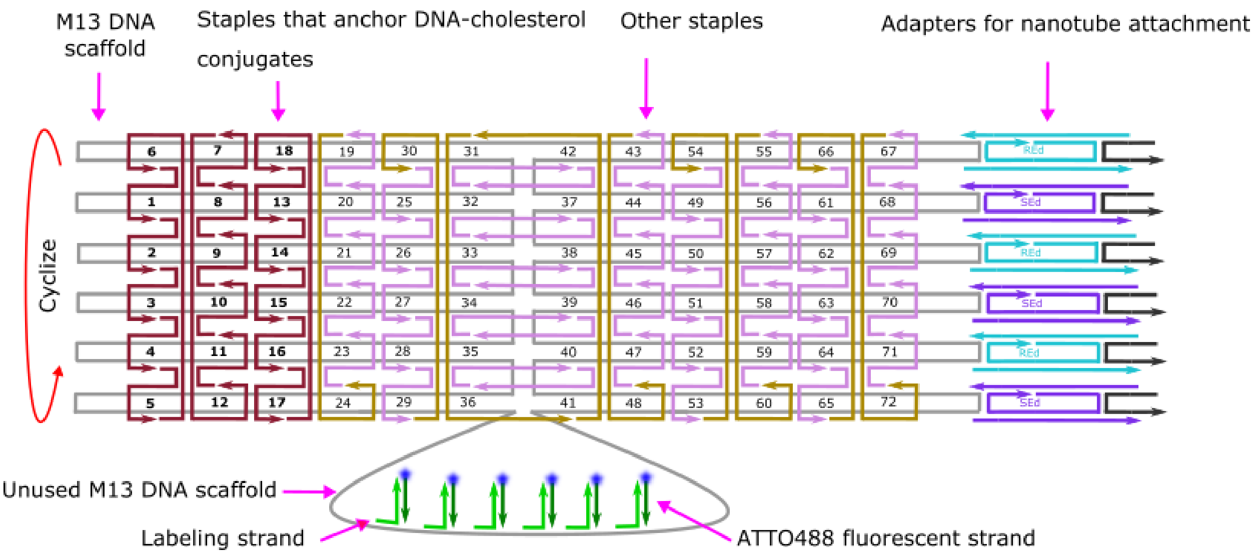


**Figure S1.** Modified Cadnano[1–3] map showing the positions of the staple strands for the DNA nanotube seed origami.The arrangement of the M13mp18 scaffold (grey) and where it interacts with the 72 ssDNA staple strands (brown and purple) of the DNA origami nanotube seed are shown. The hairpins of the first 18 ssDNA staples (lavender) were modified to present a sequence complementaryto the DNAsequence on the DNA-cholesterol conjugates used to create a hydrophobic region on the pore, allowing it to insert into the membrane. The hairpin domains of staples and domains that are binding sites for the cholesterol-DNA conjugates on the staples are not shown on the strands in the map. The adapter strands that serve as templates for nanotube growth (cyan, purple and black) are positioned on the right in the schematic. The DNA origami nanotube seed adapter monomer sequences were taken from Mohammed *et* al 2017, Figure S3.[4]

| | | | | | | | | | | | |
|---|---|---|---|---|---|---|---|---|---|---|---|
| T_5R2F_CholDNA.Tag1 | T_5R2E_CholDNA.Tag7 | T_3R2F_CholDNA.Tag13 | T_3R2E_HP | T_1R2F_HP | T_1R2E_HP | T1R2F_HP | T1R2E_HP | T3R2F_HP | T3R2E_HP | T5R2F_HP | T5R2E_HP |
| T_5R4F_CholDNA.Tag2 | T_5R4E_CholDNA.Tag8 | T_3R4F_CholDNA.Tag14 | T_3R4E_HP | T_1R4F_HP | T_1R4E_HP | T1R4F_HP | T1R4E_HP | T3R4F_HP | T3R4E_HP | T5R4F_HP | T5R4E_HP |
| T_5R6F_CholDNA.Tag3 | T_5R6E_CholDNA.Tag9 | T_3R6F_CholDNA.Tag15 | T_3R6E_HP | T_1R6F_HP | T_1R6E_HP | T1R6F_HP | T1R6E_HP | T3R6F_HP | T3R6E_HP | T5R6F_HP | T5R6E_HP |
| T_5R8F_CholDNA.Tag4 | T_5R8E_CholDNA.Tag10 | T_3R8F_CholDNA.Tag116 | T_3R8E_HP | T_1R8F_HP | T_1R8E_HP | T1R8F_HP | T1R8E_HP | T3R8F_HP | T3R8E_HP | T5R8F_HP | T5R8E_HP |
| T_5R10F_CholDNA.Tag5 | T_5R10E_CholDNA.Tag11 | T_3R10F_CholDNA.Tag17 | T_3R10E_HP | T_1R10F_HP | T_1R10E_HP | T1R10F_HP | T1R10E_HP | T3R10F_HP | T3R10E_HP | T5R10F_HP | T5R10E_HP |
| T_5R12F_CYC_CholDNA.Tag6 | T_5R12E_CYC_CholDNA.Tag12 | T_3R12F_CYC_CholDNA.Tag18 | T_3R12E_CYC_HP | T_1R12_CYC_HP | T_1R12E_CYC_HP | T_1R12F_CYC_HP | T1R12E_CYC_HP | T3R12F_CYC_HP | T3R12E_CYC_HP | T5R12F_CYC_HP | T5R12E_CYC_HP |

**Figure S2.** Schematic showing the names of the staple strands in their positions respective to the strand map of Figure S1.

The seed design was adopted from a DNA origami tall rectangle design previously published by Mohammed *et al*.[5] In the following 18 staple strands, the sequences were modified for the incorporation of cholesterol anchors.

| | | |
|---|---|---|
| T_5R2F_CholDNA.Tag1 | T_5R2E_CholDNA.Tag7 | T_3R2F_CholDNA.Tag13 |
| T_5R4F_CholDNA.Tag2 | T_5R4E_CholDNA.Tag8 | T_3R4F_CholDNA.Tag14 |
| T_5R6F_CholDNA.Tag3 | T_5R6E_CholDNA.Tag9 | T_3R6F_CholDNA.Tag15 |
| T_5R8F_CholDNA.Tag4 | T_5R8E_CholDNA.Tag10 | T_3R8F_CholDNA.Tag116 |
| T_5R10F_CholDNA.Tag5 | T_5R10E_CholDNA.Tag11 | T_3R10F_CholDNA.Tag17 |
| T_5R12F_CYC_CholDNA.Tag6 | T_5R12E_CYC_CholDNA.Tag12 | T_3R12F_CYC_CholDNA.Tag18 |

**Note S1.** DNA seed staple sequences (yellow shading denotes hairpin sequences).

| | |
|---|---|
| T_5R2F_CholDNA.Tag1 | TTGCAGCTACGTCTATGAGTTTCAAAGGAACGTCCACCGTTTTCGGTGGACTTA |
| T_5R4F_CholDNA.Tag2 | TTGCAGCTACGTCTAAAAAAAGGCTTTTGCGGTGGTCCGTTTTCGGACCACTTG |
| T_5R6F_CholDNA.Tag3 | TTGCAGCTACGTCTAACGGCTACAAGTACAACTCGGCACTTTTGTGCCGAGTTC |
| T_5R8F_CholDNA.Tag4 | TTGCAGCTACGTCTAGCTCCATGACGTAACACGGATCGCTTTTGCGATCCGTTA |
| T_5R10F_CholDNA.Tag5 | TTGCAGCTACGTCTAACGAGTAGATCAGTTGCACCGCTGTTTTCAGCGGTG TTA |
| T_5R12F_CYC_CholDNA.Tag6 | TTGCAGCTACGTCTAGGAATTACCACCACCCGTGAGGCGTTTTCGCCTCACTTT |
| T_5R2E_CholDNA.Tag7 | TTGCAGCTACGTCTAGAGAATAGGTCACCAGCGGAACCGTTTTCGGTTCCGTTT |
| T_5R4E_CholDNA.Tag8 | TTGCAGCTACGTCTAAAAGGCCGCTCCAAAACCGTGGCGTTTTCGCCACGGTTG |
| T_5R6E_CholDNA.Tag9 | TTGCAGCTACGTCTAGCGAAACAAGAGGCTTGTGCTGCGTTTTCGCAGCACTTT |
| T_5R8E_CholDNA.Tag10 | TTGCAGCTACGTCTACCAAATCATTACTTAGACGCTGGCTTTTGCCAGCGTTTC |
| T_5R10E_CholDNA.Tag11 | TTGCAGCTACGTCTAAAAGATTCTAAATTGGCGACGGACTTTTGTCCGTCGTTG |
| T_5R12E_CYC_CholDNA.Tag12 | TTGCAGCTACGTCTACTCAGAGCGAGGCATAGGCTCCGCTTTTGCGGAGCCTTG |
| T_3R2F_CholDNA.Tag13 | TGTAGCATAACTTTCAGGCATCCGTTTTCGGATGCCTTACAGTTTCTAATTGTA |
| T_3R4F_CholDNA.Tag14 | TTGCAGCTACGTCTAAGGACTGCTTACATTATTAACACTATCATAACCCACCGCCACCTCGGCTG |
| T_3R6F_CholDNA.Tag15 | TTGCAGCTACGTCTAAGGTGTGGTTACAGTTTCTAATTGTATCGGTTTAGGTCGCTGGCGACATG |
| T_3R8F_CholDNA.Tag16 | TTGCAGCTACGTCTACATGTCGCTTAGGCTTGCAAAGACTTTTTCATGATGACCCCCGAACGATG |
| T_3R10F_CholDNA.Tag17 | TTGCAGCTACGTCTACATCGTTCTTAGCGATTAAGGCGCAGACGGTCAATGACAAGAGCCTCACA |
| T_3R12F_CYC_CholDNA.Tag18 | TTGCAGCTACGTCTATGTGAGGCTTACCGGATATGGTTTAATTTCAACTACGGAACAGCAGTCCT |
| T_3R12E_CYC_HP | CCCTCAGATCGTTTACCGCTTGCGTTTTCGCAAGCGTTCAGACGACTTAATAAA |
| T_1R12F_CYC_HP | CCAAAATATACTCAGGTGCGGTCGTTTTCGACCGCATTAGGTTTAGATAGTTAG |
| T1R12E_CYC_HP | AGGGTTGAACGCTAACGCCAGGACTTTTGTCCTGGCTTGAGCGTCTGAACACCC |
| T1R12F_CYC_HP | TCTTACCATATAAGTACCGAGGCGTTTTCGCCTCGGTTTAGCCCGGAATAGGTG |
| T_1R12E_CYC_HP | TATCACCGGCGAGAGGCTGCGTCGTTTTCGACGCAGTTCTTTTGCAATCCTGAA |
| T3R12F_CYC_HP | CCTAATTTACCAGGCGTCGGAGCGTTTTCGCTCCGATTGATAAGTGGGGGTCAG |
| T3R12E_CYC_HP | TGCTCAGTGCCAGTTAGGTGGTCGTTTTCGACCACCTTCAAAATAAACAGGGAA |
| T5R12F_CYC_HP | ATTATTTAGAAGGATTGCCATCGCTTTTGCGATGGCTTAGGATTAGAAACAGTT |

| | |
|---|---|
| T5R12E_CYC_HP | CCTCAAGATCCCAATCCGTGGAGCTTTTGCTCCACGTTCAAATAAGATAGCAGC |
| T_3R2E_HP | TGCTAAACTCCACAGAGCCAGTGCTTTTGCACTGGCTTCAGCCCTCTACCGCCA |
| T_3R4E_HP | ATATATTCTCAGCTTGCCGTCCGCTTTTGCGGACGGTTCTTTCGAGTGGGATTT |
| T_3R6E_HP | CTCATCTTGGAAGTTTCGGATGGCTTTTGCCATCCGTTCCATTAAACATAACCG |
| T_3R8E_HP | AGTAATCTTCATAAGGTCTGGTCGTTTTCGACCAGATTGAACCGAACTAAAACA |
| T_3R10E_HP | ACGAACTATTAATCATGGCACCTGTTTTCAGGTGCCTTTGTGAATTTCATCAAG |
| T_1R2F_HP | CGTAACGAAAATGAATCCTGCCTGTTTTCAGGCAGGTTTTTCTGTAGTGAATTT |
| T_1R4F_HP | CTTAAACAACAACCATCGGTGCCGTTTTCGGCACCGTTCGCCCACGCGGGTAAA |
| T_1R6F_HP | ATACGTAAGAGGCAAACTCGGTCGTTTTCGACCGAGTTAGAATACACTGACCAA |
| T_1R8F_HP | CTTTGAAAATAGGCTGCCGAGGACTTTTGTCCTCGGTTGCTGACCTACCTTATG |
| T_1R10F_HP | CGATTTTAGGAAGAAACGGCAGGCTTTTGCCTGCCGTTAATCTACGGATAAAAA |
| T_1R2E_HP | ACGTTAGTTCTAAAGTCGCTTGGCTTTTGCCAAGCGTTTTTGTCGTGATACAGG |
| T_1R4E_HP | CAATGACAGCTTGATATGGCGAGCTTTTGCTCGCCATTCCGATAGTCTCCCTCA |
| T_1R6E_HP | AAACGAAATGCCACTACCACCTCGTTTTCGAGGTGGTTCGAAGGCAGCCAGCAA |
| T_1R8E_HP | CCAGGCGCGAGGACAGCTCTGGACTTTTGTCCAGAGTTATGAACGGGTAGAAAA |
| T_1R10E_HP | GGACGTTGAGAACTGGCGAGGCACTTTTGTGCCTCGTTCTCATTATGCGCTAAT |
| T1R2F_HP | AGTGTACTATACATGGCTCCTGCGTTTTCGCAGGAGTTCTTTTGATCTTTCCAG |
| T1R4F_HP | GAGCCGCCCCACCACCGTCAGGCGTTTTCGCCTGACTTGGAACCGCTGCGCCGA |
| T1R6F_HP | AATCACCACCATTTGGCGTCCTGCTTTTGCAGGACGTTGAATTAGACCAACCTA |
| T1R8F_HP | TACATACACAGTATGTCGGACCTGTTTTCAGGTCCGTTTAGCAAACTGTACAGA |
| T1R10F_HP | ATCAGAGAGTCAGAGGCGAGGTCGTTTTCGACCTCGTTGTAATTGAACCAGTCA |
| T1R2E_HP | TAAGCGTCGGTAATAACAGGAGCGTTTTCGCTCCTGTTGTTTTAACCCGTCGAG |
| T1R4E_HP | AACCAGAGACCCTCAGGCAGTCGCTTTTGCGACTGCTTAACCGCCACGTTCCAG |
| T1R6E_HP | GACTTGAGGTAGCACCGTCTGGCGTTTTCGCCAGACTTATTACCATATCACCGG |
| T1R8E_HP | TTATTACGTAAAGGTGTGGCTGCGTTTTCGCAGCCATTGCAACATACCGTCACC |
| T1R10E_HP | TGAACAAAGATAACCCAGTGCCTGTTTTCAGGCACTTTACAAGAATAAGACTCC |
| T3R2F_HP | TGCCTTGACAGTCTCTGTCGGTGCTTTTGCACCGACTTGAATTTACCCCTCAGA |
| T3R4F_HP | GCCACCACTCTTTTCACGGTCGGCTTTTGCCGACCGTTTAATCAAAT |

| | |
|---|---|
| | AGCAAGG |
| T3R6F_HP | CCGGAAACTAAAGGTGGACCTGGCTTTTGCCAGGTCTTAATTATCATAAAAGAA |
| T3R8F_HP | ACGCAAAGAAGAACTGTCGGCTCGTTTTCGAGCCGATTGCATGATTTGAGTTAA |
| T3R10F_HP | GCCCAATAGACGGGAGCACAGGCGTTTTCGCCTGTGTTAATTAACTTTCCAGAG |
| T3R2E_HP | GGAAAGCGGTAACAGTGTGGCAGCTTTTGCTGCCACTTGCCCGTATCGGGGTTT |
| T3R4E_HP | GTTTGCCACCTCAGAGACCAGGCGTTTTCGCCTGGTTTCCGCCACCGCCAGAAT |
| T3R6E_HP | TTATTCATGTCACCAAGCTCGCTGTTTTCAGCGAGCTTTGAAACCATTATTAGC |
| T3R8E_HP | ATACCCAAACACCACGCCTACCGCTTTTGCGGTAGGTTGAATAAGTGACGGAAA |
| T3R10E_HP | GCGCATTAATAAGAGCCTGGACGCTTTTGCGTCCAGTTAAGAAACAATAACGGA |
| T5R2F_HP | AATGCCCCATAAATCCGCTCGGACTTTTGTCCGAGCTTTCATTAAAAGAACCAC |
| T5R4F_HP | CACCAGAGTTCGGTCAGCCGAGCGTTTTCGCTCGGCTTTAGCCCCCTCGATAGC |
| T5R6F_HP | AGCACCGTAGGGAAGGTCGGAGGCTTTTGCCTCCGATTTAAATATTTTATTTTG |
| T5R8F_HP | TCACAATCCCGAGGAACTGGTGGCTTTTGCCACCAGTTACGCAATAATGAAATA |
| T5R10F_HP | GCAATAGCAGAGAATACCGCAGGCTTTTGCCTGCGGTTACATAAAAACAGCCAT |
| T5R2E_HP | ACAAACAACTGCCTATCACGACGCTTTTGCGTCGTGTTTTCGGAACCTGAGACT |
| T5R4E_HP | TCGGCATTCCGCCGCCGTCGCTGCTTTTGCAGCGACTTAGCATTGATGATATTC |
| T5R6E_HP | ATTGAGGGAATCAGTACGGAGCACTTTTGTGCTCCGTTGCGACAGACGTTTTCA |
| T5R8E_HP | GAAGGAAAAATAGAAAGCCTAGCGTTTTCGCTAGGCTTATTCATATTTCAACCG |
| T5R10E_HP | CTTTACAGTATCTTACCGCTCGTGTTTTCACGAGCGTTCGAAGCCCAGTTACCA |

Sequence of the 3'nanotube seed_Chol.DNAstrand:

TAGACGTAGCTGCAA/3CholTEG/.

**Note S2.** 5bp REd_SEd nanotube monomer sequences.

Figure S3 shows the schematics of the two monomers that were used to assemble DNA nanotubes. The two monomer types were termed REd and SEd respectively. The black triangles indicate the locations of the crossover points. The central strands of both REd and SEd were labeled with Cy3 fluorophores on their 5' ends. These labels made it possible to characterize the structure and lengths of nanotubes via fluorescence microscopy.

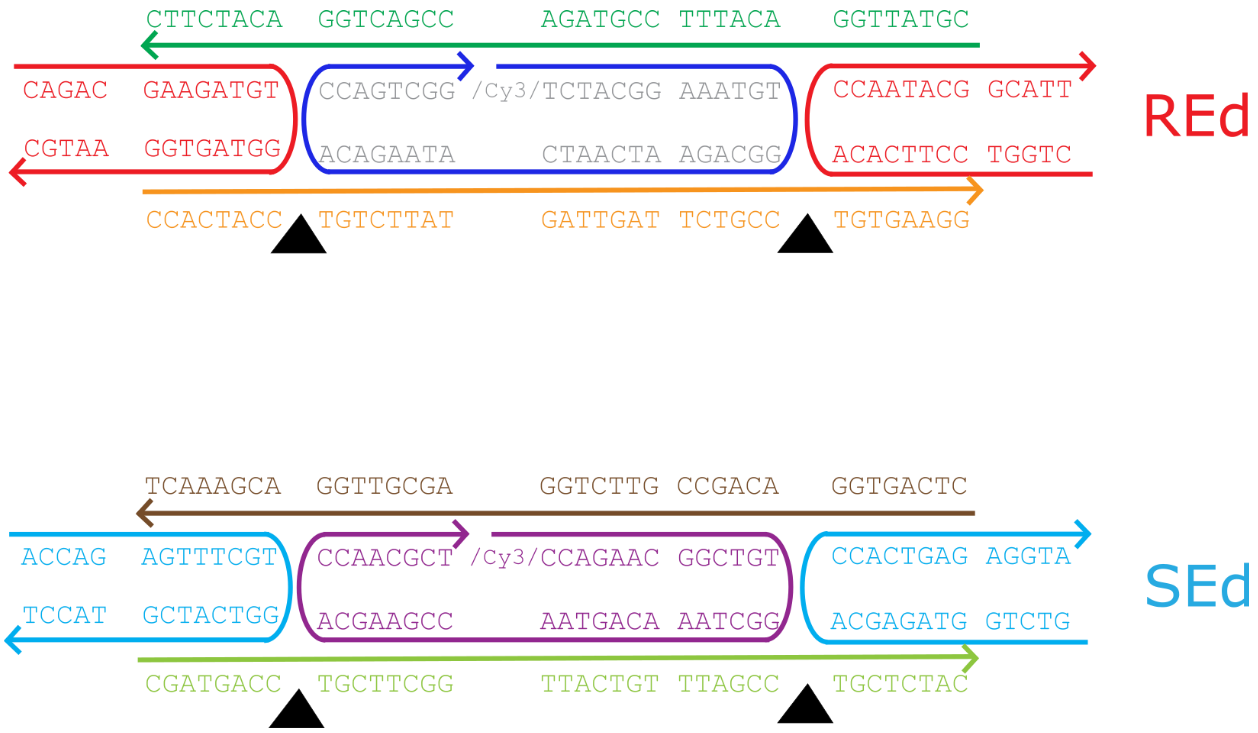


**Figure S3.** Schematic representation of the two monomers that self assemble to form the DNA nanotubes used in this work.

**<u>REd monomer strand sequences</u>**

| | |
|---|---|
| REd-strand 1 | CGTATTGGACATTTCCGTAGACCGACTGGACATCTTC |
| REd-strand 2 | CTGGTCCTTCACACCAATACGGCATT |
| REd-strand 3 | /Cy3/TCTACGGAAATGTGGCAGAATCAATCATAAGACACCAGTCGG |
| REd-strand 4 | CAGACGAAGATGTGGTAGTGGAATGC |

| | |
|---|---|
| REd-strand 5 | CCACTACCTGTCTTATGATTGATTCTGCCTGTGAAGG |

**SEd monomer strand sequences**

| | |
|---|---|
| SEd-strand 1 | CTCAGTGGACAGCCGTTCTGGAGCGTTGGACGAAACT |
| SEd-strand 2 | GTCTGGTAGAGCACCACTGAGAGGTA |
| SEd-strand 3 | /Cy3/CCAGAACGGCTGTGGCTAAACAGTAACCGAAGCACCAACGCT |
| SEd-strand 4 | ACCAGAGTTTCGTGGTCATCGTACCT |
| SEd-strand 5 | CGATGACCTGCTTCGGTTACTGTTTAGCCTGCTCTAC |

The two DNA monomer system REd and SEd used in the current study were those in Mohammed *et al* 2013, and their sequences are presented in Note S1 in that work.[5] The two monomers assembled diagonally striped lattices. We used 40 nM concentrations of the DNA strands to form nanotubes as presented in Mohammed *et al* 2013 Note S1.[5]

**Note S3.** DNA origami channel cap staple sequences.

| | |
|---|---|
| Channel cap DNA-1 | CCAAATCATTACTTAGACGCTGGCTTTTGCCAGCGTTTCCGGAACGAGGGAGTT |
| Channel cap DNA-2 | AGTAATCTTCATAAGGTCTGGTCGTTTTCGACCAGATTGAACCGAACATAACCG |
| Channel cap DNA-3 | AAAGGCCGCTCCAAAACCGTGGCGTTTTCGCCACGGTTGGAGCCTTTTCATTAC |
| Channel cap DNA-4 | CCCTCAGATCGTTTACCGCTTGCGTTTTCGCAAGCGTTCAGACGACTACCGCCA |
| Channel cap DNA-5 | GCGAAACAAGAGGCTTGTGCTGCGTTTTCGCAGCACTTTGAGGACTTACCAAGC |
| Channel cap DNA-6 | CTCATCTTGGAAGTTTCGGATGGCTTTTGCCATCCGTTCCATTAAACTAAAACA |
| Channel cap DNA-7 | TGCTAAACTCCACAGAGCCAGTGCTTTTGCACTGGCTTCAGCCCTCTTAATAAA |
| Channel cap DNA-8 | ACGAACTATTAATCATGGCACCTGTTTTCAGGTGCCTTTGTGAATTTGGGATTT |
| Channel cap DNA-9 | GAGAATAGGTCACCAGCGGAACCGTTTTCGGTTCCGTTTACAAACTACAGGTAG |
| Channel cap DNA-10 | ATATATTCTCAGCTTGCCGTCCGCTTTTGCGGACGGTTCTTTCGAGTCATCAAG |
| Channel cap DNA-11 | CTCAGAGCGAGGCATAGGCTCCGCTTTTGCGGAGCCTTGTAAGAGCCCGCCACC |
| Channel cap DNA-12 | AAAGATTCTAAATTGGCGACGGACTTTTGTCCGTCGTTGCTTGAGAAGCGGAGT |
| Channel cap DNA-13 | GAGAATAGGTCACCAGCGGAACCGTTTTCGGTTCCGTTTACAAACTCCGCCACC |
| Channel cap DNA-14 | AAAGGCCGCTCCAAAACCGTGGCGTTTTCGCCACGGTTGGAGCCTTAGCGGAGT |
| Channel cap DNA-15 | GCGAAACAAGAGGCTTGTGCTGCGTTTTCGCAGCACTTTGAGGACTAGGGAGTT |
| Channel cap DNA-16 | CCAAATCATTACTTAGACGCTGGCTTTTGCCAGCGTTTCCGGAACGTACCAAGC |
| Channel cap DNA-17 | AAAGATTCTAAATTGGCGACGGACTTTTGTCCGTCGTTGCTTGAGATTCATTAC |
| Channel cap DNA-18 | CTCAGAGCGAGGCATAGGCTCCGCTTTTGCGGAGCCTTGTAAGAGCACAGGTAG |
| Channel cap DNA-19 | TGTAGCATAACTTTCAGGCATCCGTTTTCGGATGCCTTACAGTTTCTAATTGTA |
| Channel cap DNA-20 | TCGGTTTAGGTCGCTGGCTGACGCTTTTGCGTCAGCTTAGGCTTGCAAAGACTT |
| Channel cap DNA-21 | TTTCATGATGACCCCCACCAGCCGTTTTCGGCTGGTTTAGCGATTAAGGCGCAG |
| Channel cap DNA-22 | ACGGTCAATGACAAGACGGAGGCGTTTTCGCCTCCGTTACCGGATATGGTTTAA |
| Channel cap DNA-23 | TTTCAACTACGGAACACTCGCTGCTTTTGCAGCGAGTTACATTATTAACACTAT |
| Channel cap DNA-24 | CATAACCCACCGCCACCTGGCTCGTTTTCGAGCCAGTTCCTCAGAAACAACGCC |
| Channel cap DNA-25 | CGTAACGAAAATGAATCCTGCCTGTTTTCAGGCAGGTTTTTCTGTAGTGAATTT |
| Channel cap DNA-26 | CTTAAACAACAACCATCGGTGCCGTTTTCGGCACCGTTCGCCCACGCGGGTAAA |

| | |
|---|---|
| Channel cap DNA-27 | ATACGTAAGAGGCAAACTCGGTCGTTTTCGACCGAGTTAGAATACACTGACCAA |
| Channel cap DNA-28 | CTTTGAAAATAGGCTGCCGAGGACTTTTGTCCTCGGTTGCTGACCTACCTTATG |
| Channel cap DNA-29 | CGATTTTAGGAAGAAACGGCAGGCTTTTGCCTGCCGTTAATCTACGGATAAAAA |
| Channel cap DNA-30 | CCAAAATATACTCAGGTGCGGTCGTTTTCGACCGCATTAGGTTTAGATAGTTAG |
| Channel cap DNA-31 | ACGTTAGTTCTAAAGTCGCTTGGCTTTTGCCAAGCGTTTTTGTCGTGATACAGG |
| Channel cap DNA-32 | CAATGACAGCTTGATATGGCGAGCTTTTGCTCGCCATTCCGATAGTCTCCCTCA |
| Channel cap DNA-33 | AAACGAAATGCCACTACCACCTCGTTTTCGAGGTGGTTCGAAGGCAGCCAGCAA |
| Channel cap DNA-34 | CCAGGCGCGAGGACAGCTCTGGACTTTTGTCCAGAGTTATGAACGGGTAGAAAA |
| Channel cap DNA-35 | GGACGTTGAGAACTGGCGAGGCACTTTTGTGCCTCGTTCTCATTATGCGCTAAT |
| Channel cap DNA-36 | TATCACCGGCGAGAGGCTGCGTCGTTTTCGACGCAGTTCTTTTGCAATCCTGAA |
| Channel cap DNA-37 | AGTGTACTATACATGGCTCCTGCGTTTTCGCAGGAGTTCTTTTGATCTTTCCAG |
| Channel cap DNA-38 | GAGCCGCCCCACCACCGTCAGGCGTTTTCGCCTGACTTGGAACCGCTGCGCCGA |
| Channel cap DNA-39 | AATCACCACCATTTGGCGTCCTGCTTTTGCAGGACGTTGAATTAGACCAACCTA |
| Channel cap DNA-40 | TACATACACAGTATGTCGGACCTGTTTTCAGGTCCGTTTAGCAAACTGTACAGA |
| Channel cap DNA-41 | ATCAGAGAGTCAGAGGCGAGGTCGTTTTCGACCTCGTTGTAATTGAACCAGTCA |
| Channel cap DNA-42 | TCTTACCATATAAGTACCGAGGCGTTTTCGCCTCGGTTTAGCCCGGAATAGGTG |
| Channel cap DNA-43 | TAAGCGTCGGTAATAACAGGAGCGTTTTCGCTCCTGTTGTTTTAACCCGTCGAG |
| Channel cap DNA-44 | AACCAGAGACCCTCAGGCAGTCGCTTTTGCGACTGCTTAACCGCCACGTTCCAG |
| Channel cap DNA-45 | GACTTGAGGTAGCACCGTCTGGCGTTTTCGCCAGACTTATTACCATATCACCGG |
| Channel cap DNA-46 | TTATTACGTAAAGGTGTGGCTGCGTTTTCGCAGCCATTGCAACATACCGTCACC |
| Channel cap DNA-47 | TGAACAAAGATAACCCAGTGCCTGTTTTCAGGCACTTTACAAGAATAAGACTCC |
| Channel cap DNA-48 | AGGGTTGAACGCTAACGCCAGGACTTTTGTCCTGGCTTGAGCGTCTGAACACCC |
| Channel cap DNA-49 | TGCCTTGACAGTCTCTGTCGGTGCTTTTGCACCGACTTGAATTTACCCCTCAGA |
| Channel cap DNA-50 | GCCACCACTCTTTTCACGGTCGGCTTTTGCCGACCGTTTAATCAAATAGCAAGG |
| Channel cap DNA-51 | CCGGAAACTAAAGGTGGACCTGGCTTTTGCCAGGTCTTAATTATCATAAAAGAA |
| Channel cap DNA-52 | ACGCAAAGAAGAACTGTCGGCTCGTTTTCGAGCCGATTGCATGATTTGAGTTAA |
| Channel cap DNA-53 | GCCCAATAGACGGGAGCACAGGCGTTTTCGCCTGTGTTAATTAACTTTCCAGAG |
| Channel cap | CCTAATTTACCAGGCGTCGGAGCGTTTTCGCTCCGATTGATAAGTGGGGGTCA |

| | |
|---|---|
| DNA-54 | G |
| Channel cap DNA-55 | GGAAAGCGGTAACAGTGTGGCAGCTTTTGCTGCCACTTGCCCGTATCGGGGTTT |
| Channel cap DNA-56 | GTTTGCCACCTCAGAGACCAGGCGTTTTCGCCTGGTTTCCGCCACCGCCAGAAT |
| Channel cap DNA-57 | TTATTCATGTCACCAAGCTCGCTGTTTTCAGCGAGCTTTGAAACCATTATTAGC |
| Channel cap DNA-58 | ATACCCAAACACCACGCCTACCGCTTTTGCGGTAGGTTGAATAAGTGACGGAAA |
| Channel cap DNA-59 | GCGCATTAATAAGAGCCTGGACGCTTTTGCGTCCAGTTAAGAAACAATAACGGA |
| Channel cap DNA-60 | TGCTCAGTGCCAGTTAGGTGGTCGTTTTCGACCACCTTCAAAATAAACAGGGAA |
| Channel cap DNA-61 | AATGCCCCATAAATCCGCTCGGACTTTTGTCCGAGCTTTCATTAAAAGAACCAC |
| Channel cap DNA-62 | CACCAGAGTTCGGTCAGCCGAGCGTTTTCGCTCGGCTTTAGCCCCCTCGATAGC |
| Channel cap DNA-63 | AGCACCGTAGGGAAGGTCGGAGGCTTTTGCCTCCGATTTAAATATTTTATTTTG |
| Channel cap DNA-64 | TCACAATCCCGAGGAACTGGTGGCTTTTGCCACCAGTTACGCAATAATGAAATA |
| Channel cap DNA-65 | GCAATAGCAGAGAATACCGCAGGCTTTTGCCTGCGGTTACATAAAAACAGCCAT |
| Channel cap DNA-66 | ATTATTTAGAAGGATTGCCATCGCTTTTGCGATGGCTTAGGATTAGAAACAGTT |
| Channel cap DNA-67 | ACAAACAACTGCCTATCACGACGCTTTTGCGTCGTGTTTTCGGAACCTGAGACT |
| Channel cap DNA-68 | TCGGCATTCCGCCGCCGTCGCTGCTTTTGCAGCGACTTAGCATTGATGATATTC |
| Channel cap DNA-69 | ATTGAGGGAATCAGTACGGAGCACTTTTGTGCTCCGTTGCGACAGACGTTTTCA |
| Channel cap DNA-70 | GAAGGAAAAATAGAAAGCCTAGCGTTTTCGCTAGGCTTATTCATATTTCAACCG |
| Channel cap DNA-71 | CTTTACAGTATCTTACCGCTCGTGTTTTCACGAGCGTTCGAAGCCCAGTTACCA |
| Channel cap DNA-72 | CCTCAAGATCCCAATCCGTGGAGCTTTTGCTCCACGTTCAAATAAGATAGCAGC |

1

**Note S4.** Stock preparations.

DNA origami nanotube seed staple mixes

The staple mix for the DNA origami nanotube seeds contained each of the staples listed in Note S1 at 3.70 μM suspended in water. The cholesterol DNA conjugate staple mixtures contained each of the staples that bind the cholesterol DNA conjugate at concentrations of 50 μM. The nanotube seed cholesterol DNA conjugate (3' nanotube seed_Chol.DNAstrand) is prepared at a concentration of 100 μM (see Note S1 for sequences).

DNA origami channel cap staple mixes

The DNA origami channel cap staple mix contained each of the staples listed in Note S3 at 2.38 μM suspended in water.

Adapter strand mixes for the DNA origami channel caps

The DNA origami channel caps adapter monomer sequences were taken from Mohammed *et al* 2017,[4] Figure S4. Each adapter strand mix for the DNA origami channel caps contained the strands Channel cap REd Ad_1, Channel cap REd Ad_2, Channel cap REd Ad_3, Channel cap SEd Ad_1, Channel cap SEd Ad_2, Channel cap SEd Ad_3, Channel cap REd Ad_4, Channel cap REd Ad_5, Channel cap REd Ad_6, Channel cap SEd Ad_4, Channel cap SEd Ad_5, Channel cap SEd Ad_6, Channel cap REd Ad_7, Channel cap REd Ad_8, Channel cap REd Ad_9, Channel cap SEd Ad_7, Channel cap SEd Ad_8 and Channel cap SEd Ad_9 at 1 μM and the strands with the sticky ends, namely Channel cap REd sticky Ad_1, Channel cap SEd sticky Ad_1,

Channel cap REd sticky Ad_2, Channel cap SEd sticky Ad_2, Channel cap REd sticky Ad_3 and Channel cap SEd sticky Ad_3 at 2 µM suspended in water.

DNA origami nanotubes seed labeling attachment strands mix

The attachment strand mix contained 1 µM of each of 100 labeling attachment strands. The attachment strand sequences were those listed in Mohammed *et al* 2017, Supplementary Information Section 1.3.[4]

Labeling strand sequences

The labeling strands used in the current study were taken from Mohammed *et al* 2017, Supplementary Information Section 1.3.[4] Their sequences are:

Labeling_strand_ATTO488: /5ATTO488N/AAGCGTAGTCGGATCTC

Labeling_strand_ATTO647N: /5ATTO647NN/AAGCGTAGTCGGATCTC

The DNA nanotube seed origami was labeled with ATTO488 (blue), DNA origami channel cap with ATTO647N (red) and the DNA nanotubes with Cy3 (green) labeling dyes respectively.

**Note S5.** DNA origami channel cap adapter monomer sequences for control measurements.

For control experiments involving the channel caps and the DNA origami nanotube seeds the 6 sticky end strands were removed from the channel cap adapters to prevent the channel caps and the seeds from binding.The nanotube seed has adapter strands that bind to the sequences on one side of the seed, while the channel cap has a different set of adapter strands that bind to the other side of the seed. Since the six sticky end strands were removed from the channel caps, the adapter strands of the channel caps will not facilitate binding to the nanotube seed. The complete list of adapter strand sequences for the nanotube seed and the channel caps can be found in Mohammed *et al* 2017, Figures S3 and S4. [4]Each adapter strand mix for the channel cap contained the adapters at 1 μM suspended in water.

Channel caps with the sticky end strands removed for the control experiments

| | |
|---|---|
| Channel cap REd sticky Ad_1 | CAGACGAGTGCAGAGTCAGCGAATGC |
| Channel cap SEd sticky Ad_1 | ACCAGACTCCATCGGTTGTGGTACCT |
| Channel cap REd sticky Ad_2 | CAGACCGTAGGCTGGCAGAGCAATGC |
| Channel cap SEd sticky Ad_2 | ACCAGCACTCTGTAGCGTGACTACCT |
| Channel cap REd sticky Ad_3 | CAGACCAGAGGTCAGTCGTGGAATGC |
| Channel cap SEd sticky Ad_3 | ACCAGCAGTGTCGCAGTCCGCTACCT |

Adapter strands for the channel caps used for the control experiments

| | |
|---|---|
| Channel cap_control_REd Ad_1 | AGGGATAGCAAGCCCACAACGTGAGGACACTTGGAGGCTGCACT C |
| Channel cap_control_REd Ad_2 | TGTCCTCACGTTGCTGGATGCCGATCCTACGACACCTCCAAG |
| Channel cap_control_REd Ad_3 | CGCTGACTTGTCGTAGGATCGGCATCCAGATAGGAACCCATGTA C |
| Channel cap_control_SEd Ad_1 | GAATTGCGAATAATAAGTGACCTTGCTGTACCGTCGAGATGGAGT |
| Channel cap_control_SEd Ad_2 | ACAGCAAGGTCACCGCAGTTGGCACTAGGCGACATCGACGGT |
| Channel cap_control_SEd Ad_3 | CCACAACCTGTCGCCTAGTGCCAACTGCGTTTTTTCACGTTGAAA |
| Channel cap_control_REd Ad_4 | ACCCTCAGCAGCGAAACGAGTACGGCAACACGGTGAGAGCCTAC G |

| | |
|---|---|
| Channel cap_control_REdAd_5 | GTTGCCGTACTCGACTGGTCACGAACGTCTCCAACTCACCGT |
| Channel cap_control_REdAd_6 | GCTCTGCCTTGGAGACGTTCGTGACCAGTGACAGCATCGGAACGA |
| Channel cap_control_SEdAd_4 | TGTATCATCGCCTGATCAACGGTACGAGATGCGAAGCACAGAGTG |
| Channel cap_control_SEdAd_5 | TCTCGTACCGTTGCCAGTAGACCTAGCCGACGTGGCTTCGCA |
| Channel cap_control_SEdAd_6 | GTCACGCTCACGTCGGCTAGGTCTACTGGAAATTGTGTCGAAATC |
| Channel cap_control_REdAd_7 | CATTCAGTGAATAAGGACGCTATGCCTATCGCTCTAGGACCTCTG |
| Channel cap_control_REdAd_8 | ATAGGCATAGCGTTGCTCCAGTCTGCTGCTCAGGCTAGAGCG |
| Channel cap_control_REdAd_9 | CCACGACTCCTGAGCAGCAGACTGGAGCACTTGCCCTGACGAGAA |
| Channel cap_control_SEdAd_7 | GAATACCACATTCAACACCGATGAGGATCACGGCACTCGACACTG |
| Channel cap_control_SEdAd_8 | GATCCTCATCGGTCAAGCGAAGGTGCGAGCCTGTAGTGCCGT |
| Channel cap_control_SEdAd_9 | GCGGACTGACAGGCTCGCACCTTCGCTTGTAATGCAGATACATAA |

**Table S1.** DNA nanotube seed assembly mixture.

| S. No | Seed assembly mixture | Working concentration [nM] | Stock concentration [nM] | Volume to be added [µl] |
|---|---|---|---|---|
| 1. | Water | – | – | 25.25 |
| 2. | 10X TAEM | 1 | 10 | 5.0 |
| 3. | BSA [mg/mL] | 0.05 | 1 | 2.50 |
| 4. | Staple strands mix | 250 | 3700 | 3.38 |
| 5. | Cholesterol conjugate staple strands 1–6 | 250 | 50000 | 0.25 |
| 6. | Cholesterol conjugate staple strands 7–12 | 250 | 50000 | 0.25 |
| 7. | Cholesterol conjugate staple strands 13–18 | 250 | 33330 | 0.38 |
| 8. | Adapter strands mix [DNA nanotube seed adapters] | 100 | 1000 | 5.0 |
| 9. | M13mp18 scaffold | 5 | 100 | 2.50 |
| 10. | Attachment strand mix | 10 | 1000 | 0.50 |
| 11. | ATTO488 labeling strand | 1000 | 10000 | 5.0 |
| | Total volume | | | 50 |

**Note S6.** Protocol for annealing and purifying DNA origami nanotube seeds.

Step 1. Total volume of 50 µL (see Table S1) containing M13mp18 scaffold, staple strands, cholesterol conjugate staple strands, adapter strands, attachment strands and the dye strands in 1X TAEM buffer (40 mM Tris-acetate, 1 mM EDTA and 12.5 mM magnesium acetate) was added to a PCR tube. BSA was included to reduce DNA adsorption to PCR tubes.

Step 2. The seeds were annealed using the protocol for annealing seeds given in Mohammed *et al*.[5] Briefly, the seed assembly mixture was kept in an Eppendorf Mastercycler in a 5-stage reaction. The seed samples were initially heated at 90°C for 5 min to melt all DNA, then annealed from 90 to 45°C at 1°C/min, and then incubated at 45°C for 1 h to allow the formation of seed origami nanotube monomers. Samples were slowly cooled from 45 to 32°C at 0.1°C/min to a final temperature of 32°C.

<u>Nanotube seed purification for lipid bilayer experiments</u>

The annealed seeds were then mixed with the cholesterol-DNA conjugate (3' nanotube seed_Chol.DNA) present at a final concentration of 10 µM. Prior to mixing the 3' nanotube seed_Chol.DNA with the seeds, the strands were heated at 50°C for 15 min. The seeds and the 3' nanotube seed_Chol.DNA strand were then incubated at room temperature for 1 h. After incubation, the nanotube seeds were centrifuged using 100-kDa MWCO Amicon filters (Amicon® ultra centrifugal filters – 100 K) and 1X TAEM buffer. To 50 µL of the nanotube seed solution, 350 µL of the 1X TAEM buffer was added and centrifuged at 3000 g for 30 min. The flow-through was discarded each time and the filter was replaced with 350 µL of the fresh 1X TAEM buffer. After 3–5 times, the filter was turned upside down into a new amicon tube and centrifuged at 3000 g for 2 min to collect the final nanotube seed solution.

**Note S7.** Protocol for growing and purifying seeded nanotubes.

Step 1. Seeded nanotube monomer mixes were prepared by first mixing 15.9 µL of Milli-Q water, 1 µL of 1 mg/mL BSA, 0.8 µL of 1000 nM 5bp REd_SEd nanotube monomer mix solution (Final concentration: 40 nM, see Note S2 and Figure S3 for sequences) and 2 µL 10X TAEM buffer to a final volume of 19.7 µL per monomer solution.

Step 2. The seeds were prepared and annealed following Note S6 and purified by centrifuging the contents at 3000 g for 4 min using 100-kDa MWCO Amicon filters (Amicon® ultra centrifugal filters – 100 K), four times. After all of the centrifugation steps, the concentration of the purified nanotube seeds was measured as described and a solution of 1.2 nM seeds was prepared by mixing more concentrated seeds with 1XTAEM buffer.[6]

Step 3. The monomer solutions from step 1 were annealed from 90°C to 32°C following the protocol given in Step 2 of Note S6.[5] When the solution reached to 32°C, 0.3 µL of the nanotube seed solution (stock concentration 1.2 nM) was added to 19.7 µL of the nanotube monomer solution so that the final seed and monomer concentration were 0.018 nM and 40 nM respectively. The reaction mix was incubated at 32°C for almost 18–24 h and then imaged using fluorescence microscopy to check the yield of seeded tubes and the tube length.

4. For the lipid bilayer experiments, 10 µM of 3' nanotube seed_Chol.DNA strand was incubated with the seeded nanotube solution for 1 h under ambient conditions. The 3' nanotube seed_Chol.DNA strand was heated at 50°C for 15 min prior mixing them with the seeded nanotube solution. After incubation, the seeded nanotube solution was centrifuged using 100-kDa MWCO Amicon filters (Amicon®ultra centrifugal filters – 100

K) and 1X TAEM buffer. To 50 µL of the nanotube seed solution, 350 µL of the 1X TAEM buffer was added and centrifuged at 3000 g for 30 min. The flow-through was discarded each time and the filter was replaced with 350 µL of the fresh 1X TAEM buffer. After 3–5 times, the filter is turned upside down into a new amicon tube and centrifuged at 3000 g for 2 min to collect the final seeded nanotube solution.

**Table S2.** Unseeded nanotube assembly mixture.

| S. No | Unseeded nanotube assembly mixture | Working concentration [nM] | | Stock concentration [nM] | Volume to be added [µL] |
|---|---|---|---|---|---|
| 1. | Water | – | | – | 30 |
| 2. | 10X TAEM | 1 | | 10 | 5 |
| 3. | BSA [mg/mL] | 0.05 | | 1 | 2.5 |
| 4. | 5bp REd_SEd monomer mix | 250 | | 1000 | 12.5 |
| | Total volume | | | | 50 |

**Note S8.** Protocol for growing unseeded nanotubes.

Step 1. Total volume of 50 µL (see Table S2) containing the above components in 1X TAEM buffer was added to a PCR tube. BSA was included to reduce DNA adsorption to PCR tubes.

Step 2. The monomer solutions from Step 1 were annealed from 90°C to 32°C following the protocol given in Step 2 of Note S6.[5] The reaction mix was further incubated at 32°C for almost 18–24 h and then imaged using fluorescence microscopy to check the yield of the unseeded nanotubes.

Step 3. For the lipid bilayer experiments, the unseeded nanotube solution was filter purified 4 times at 3000 g for 4 min using 100-kDa MWCO Amicon filters (Amicon® ultra centrifugal filters – 100 K) and 1X TAEM buffer. To 50 µL of the unseeded nanotube solution, 350 µL of the 1X TAEM buffer was added and centrifuged at 3000 g for 4 min. The flow-through was discarded each time and the filter was replaced with 350 µL of the fresh 1X TAEM buffer. After 4 times, the filter was turned upside down into a new Amicon tube and centrifuged at 3000 g for 2 min to collect the final unseeded nanotube solution.

**Table S3.** DNA origami channel cap assembly mixture.

| S. No | Channel cap assembly mixture | Working concentration [nM] | Stock concentration [nM] | Volume to be added [µL] |
|---|---|---|---|---|
| 1. | Water | – | – | 24.25 |
| 2. | 10X TAEM | 1 | 10 | 5.0 |
| 3. | BSA [mg/mL] | 0.05 | 1 | 2.50 |
| 4. | Channel cap staple strands mix | 250 | 2380 | 5.25 |
| 5. | Adapter strands mix | 100 | 1000 | 5.0 |
| 6. | M13mp18 scaffold | 5 | 100 | 2.50 |
| 7. | Attachment strand mix | 10 | 1000 | 0.50 |
| 8. | ATTO647N labeling strand | 1000 | 10000 | 5.0 |
| | Total volume | | | 50 |

**Note S9.** Protocol for assembling and purifying channel caps and nanotube seed-channel cap complexes.

Step 1. Total volume of 50 µL (see Table S3) containing M13mp18 scaffold, channel cap staple strands, adapter strands, attachment strands and dye strand in 1X TAEM buffer was added to a PCR tube. BSA was included to reduce the adsorption of DNA to PCR tubes.

Step 2. The channel caps and the nanotube seeds were prepared and annealed using the protocol as given in Note S6.[5]

Step 3. The assembled channel caps and the nanotube seeds were filter purified four times at 3000 g for 4 min using 100-kDa MWCO Amicon filters (Amicon® ultra centrifugal filters – 100 K) and 1X TAEM buffer. To 50 µL of the channel cap or the nanotube seed solution, 350 µL of the 1X TAEM buffer was added and centrifuged at 3000 g for 4 min. The flow-through was discarded each time and the filter was replaced with 350 µL of the fresh 1X TAEM buffer. After 4 times, the filter was turned upside down into a new amicon tube and centrifuged at 3000 g for 2 min to collect the final channel cap or the nanotube seed solution. The structures were then visualized under transmission electron microscopy.

Step 4. For the nanotube seed-channel cap complex mix, the purified channel caps were added to the purified seed solution 3-fold in excess to make sure all the nanotube seeds have channel caps. The channel capped nanotube seeds solution was incubated for 3 h at 32°C and finally incubated with the pre-heated cholesterol-DNA conjugate (3' nanotube seed_Chol.DNA) for 1 h under room temperature.

Step 5. The final channel capped nanotube seed solution was purified via spin filtration at 3000 g for 30 min using 100-kDa MWCO Amicon filters (Amicon® ultra centrifugal filters – 100 K) and 1X TAEM buffer. To 50 µL of the channel capped nanotube seed complex solution, 350 µL of the 1X TAEM buffer was added and centrifuged at 3000 g for 30 min. The flow-through was discarded each time and the filter was replaced with 350 µL of the fresh 1X TAEM buffer. After 2–3 times, the filter is turned upside down into a new amicon tube and centrifuged at 3000 g for 2 min to collect the final nanotube seed-channel cap complex solution. The channel capped nanotube seeds were then visualized under fluorescence and transmission electron microscopy to measure the yield and the folding nature of the seed-channel capped complexes.

**Note S10.** Protocol for assembling and purifying seeded nanotube-channel cap complexes.

Step 1. The seeded nanotubes and the channel caps were annealed and purified as described in Notes S7 and S9. The folded channel caps were then added at 3-fold excess to the seeded nanotubes to make sure all the nanotubes had channel caps. The channel capped nanotube seed solution was incubated for 3 h at 32°C and finally incubated with the pre-heated DNA-cholesterol conjugate (3' nanotube seed_Chol.DNA) for 1 h under room temperature. The final channel capped seeded nanotube solution was further purified via spin filtration at 3000 g for 30 min using 100-kDa MWCO Amicon filters (Amicon® ultra centrifugal filters – 100 K) and 1X TAEM buffer. To 50 µL of the channel capped seeded nanotube complex solution, 350 µL of the 1X TAEM buffer was added and centrifuged at 3000 g for 30 min. The flow-through was discarded each time and 350 µL of the fresh 1X TAEM buffer was added back to the filter. After 2–3 cycles of this process, the filter was turned upside down into a new amicon tube and centrifuged at 3000 g for 2 min to collect the final seeded nanotube-channel cap complex solution. The channel capped seeded nanotubes were then visualized under fluorescence microscopy to measure the yield of the channel capped seeded tube complexes.

**Table S4.** Channel cap assembly mixture for control measurements where adapters mediating channel cap-seed binding are omitted.

| S. No | Channel cap assembly mixture without the sticky end adapter strands | Working concentration [nM] | Stock concentration [nM] | Volume to be added [µL] |
|---|---|---|---|---|
| 1. | Water | – | – | 24.25 |
| 2. | 10X TAEM | 1 | 10 | 5.0 |
| 3. | BSA [mg/mL] | 0.05 | 1 | 2.50 |
| 4. | Channel cap staple strands mix | 250 | 2380 | 5.25 |
| 5. | Adapter strands mix (sticky end strands removed (See Note S5) | 100 | 1000 | 5.0 |
| 6. | M13mp18 scaffold | 5 | 100 | 2.50 |
| 7. | Attachment strand mix | 10 | 1000 | 0.50 |
| 8. | ATTO647N labeling strand | 1000 | 10000 | 5.0 |
| | Total volume | | | 50 |

**Note S11.** Protocol for growing channel caps without sticky end adapters.

Step 1. Total volume of 50 µL (see Table S4) containing M13mp18 scaffold, channel cap staple strands, adapter strands (without the sticky ends, See Note S5), attachment strands and dye strand in 1X TAEM buffer was added to a PCR tube. BSA was included to reduce DNA adsorption to PCR tubes.

Step 2. The nanotube seeds and the channel caps were annealed using the protocol given in Note S6.[5]

Step 3. The assembled channel caps and the nanotubes seeds were filter purified four times at 3000 g for 4 min using 100-kDa MWCO Amicon filters (Amicon® ultra centrifugal filters – 100 K) and 1X TAEM buffer.

Step 4. For the nanotube seed-channel cap complex mix, the purified channel caps were added to the nanotube seed solution 3-fold in excess to make sure all the nanotube seeds have channel caps. The channel capped nanotube seed solution was incubated for 3 h at 32°C and finally incubated with the pre-heated cholesterol-DNA conjugate (3' nanotube seed_Chol.DNA) for 1 h under room temperature.

Step 5. The final channel capped nanotube seed solution was purified via spin filtration at 3000 g for 30 min using 100-kDa MWCO Amicon filters (Amicon® ultra centrifugal filters – 100 K) and 1X TAEM buffer. To 50 µL of the channel capped nanotube seed complex solution, 350 µL of the 1X TAEM buffer was added and centrifuged at 3000 g for 30 min. The flow-through was discarded each time and the filter was replaced with 350 µL of the fresh 1X TAEM buffer. After 2–3 times, the filter is turned upside down into a new amicon tube and centrifuged at 3000 g for 2 min to collect the final solution.

The resulting solution was then visualized under fluorescence microscopy to ensure that there is not binding between the seeds and the channel caps.

**Note S12.** A theoretical model of channel conductance.

Assuming the DNA seed origami is an ideal cylinder and that the passage of ions in solution through the channel is the sole mediator of conductance, a simple estimate for the conductance of a DNA seed channel could be obtained using its dimensions (Length = 63 nm and inner diameter = 7.3 nm). The conductance of such as cylindrical pore is given by the equation[7]

$$G = \frac{\pi d^2}{4\mathrm{L} + \pi d} K$$

where, *d* is the diameter of the seed origami; *L* is length of the DNA origami channel and *K* is the molar conductivity of 1 M KCl, 10 mM $MgCl_2$ at 23°C, *i.e.* *K* = 8 S/m. Using these values, the conductance predicted by this theoretical model is 4.8 nS. As noted in the main text, this model does not appear to be predictive of the conductance of synthetic DNA membrane channels, and this prediction does not match our measured conductance (on order 1–1.5 nS).

This model also notably, predicts that conductance should decrease with channel length. Thus, for a nanotube with length 1 µm, this formula would predict a conductance of 0.33 nS, which is much smaller than the observed conductance of on order 1 nS. The lack of dependence of channel length on conductance further confirms that this model, which assumes that ions transit from channel end to channel end, is not a good predictor of conductance of the origami seed or nanotube seed channel.

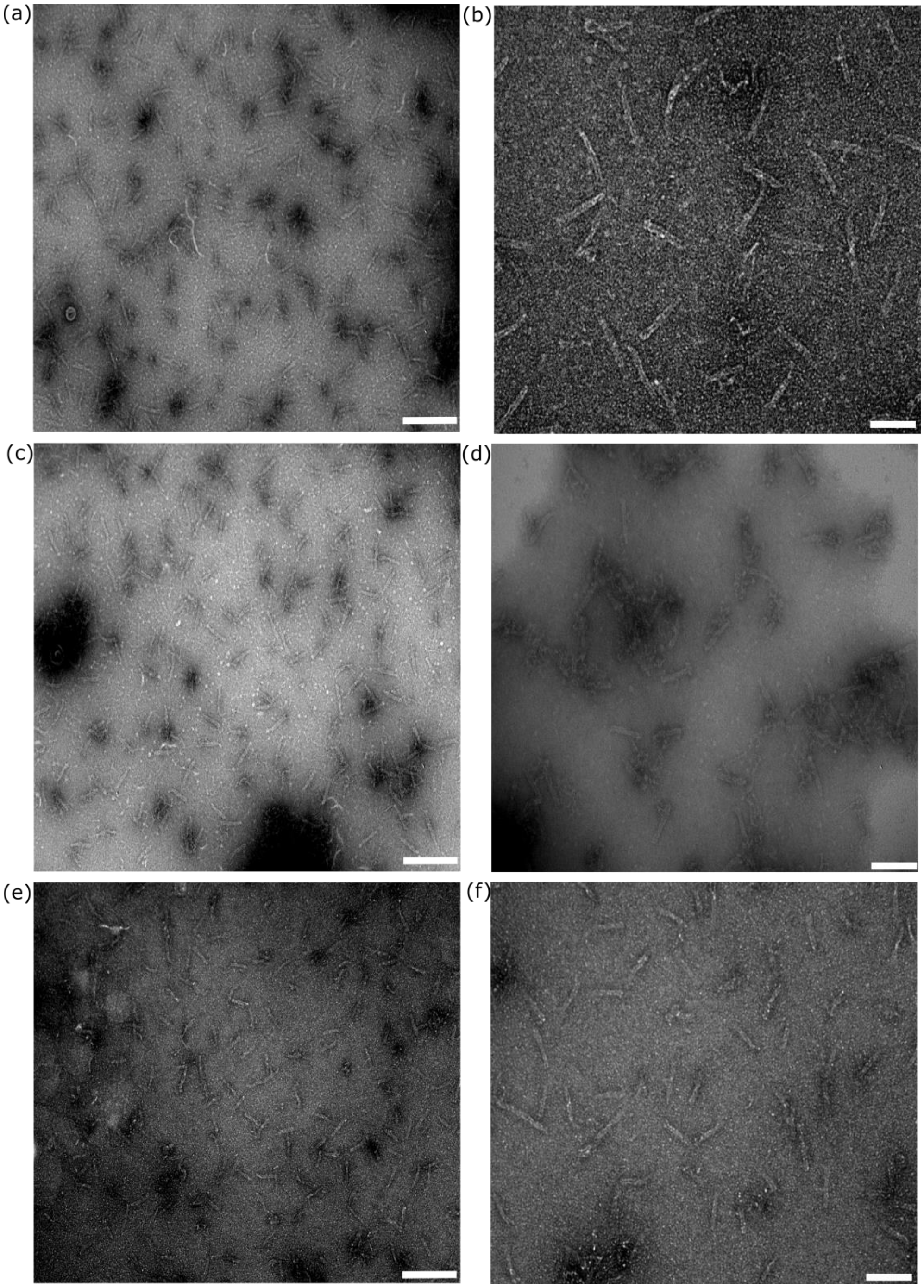


**Figure S4.** Negative stain transmission electron micrographs of DNA origami nanotube seeds. (**a**, **b**) DNA origami nanotube seeds in 1X TAEM buffer in the absence of cholesterol-DNA conjugates. The structures were centrifuged using 100-kDa MWCO Amicon filters (Amicon® ultra centrifugal filters – 100 K) and 1X TAEM buffer. To 50 µL of the nanotube seed solution, 350 µL of the 1X TAEM buffer was added and centrifuged at 3000 g for 4 min. The flow-through was discarded each time and the filter was replaced with 350 µL of the fresh 1X TAEM buffer. After four times, the filter was turned upside down into a new amicon tube and centrifuged at 3000 g for 2 min to collect the final nanotube seed solution; (**c**, **d**) DNA origami seeds in 1X TAEM buffer, after filter purification (3000 g for 4 min, 4 times) in the presence of cholesterol-DNA conjugates (3' nanotube seed_Chol.DNA); and (**e**, **f**) DNA origami seeds devoid of cholesterol-DNA conjugates in the electrophysiology buffer: 1 M KCl, 10 mM $Mg^{2+}$. The nanotube seeds were annealed using the same protocol as shown in Note S6. After annealing, the structures were filter purified using 100-kDa amicon filters (Amicon® ultra centrifugal filters – 100 K) and 1 M KCl, 10 mM $MgCl_2$. To 50 µL of the nanotube seed solution, 350 µL of the 1 M KCl, 10 mM $MgCl_2$ buffer was added and centrifuged at 3000 g for 4 min. The flow-through was discarded each time and the filter was replaced with 350 µL of the fresh 1 M KCl, 10 mM $MgCl_2$ buffer. After 4 times, the filter was turned upside down into a new amicon tube and centrifuged at 3000 g for 2 min to collect the final nanotube seed solution. 2% uranyl acetate was used as the staining agent. Scale bars: **a**, **c**, **e**: 200 nm; **b**, **d** and **f**: 100 nm.

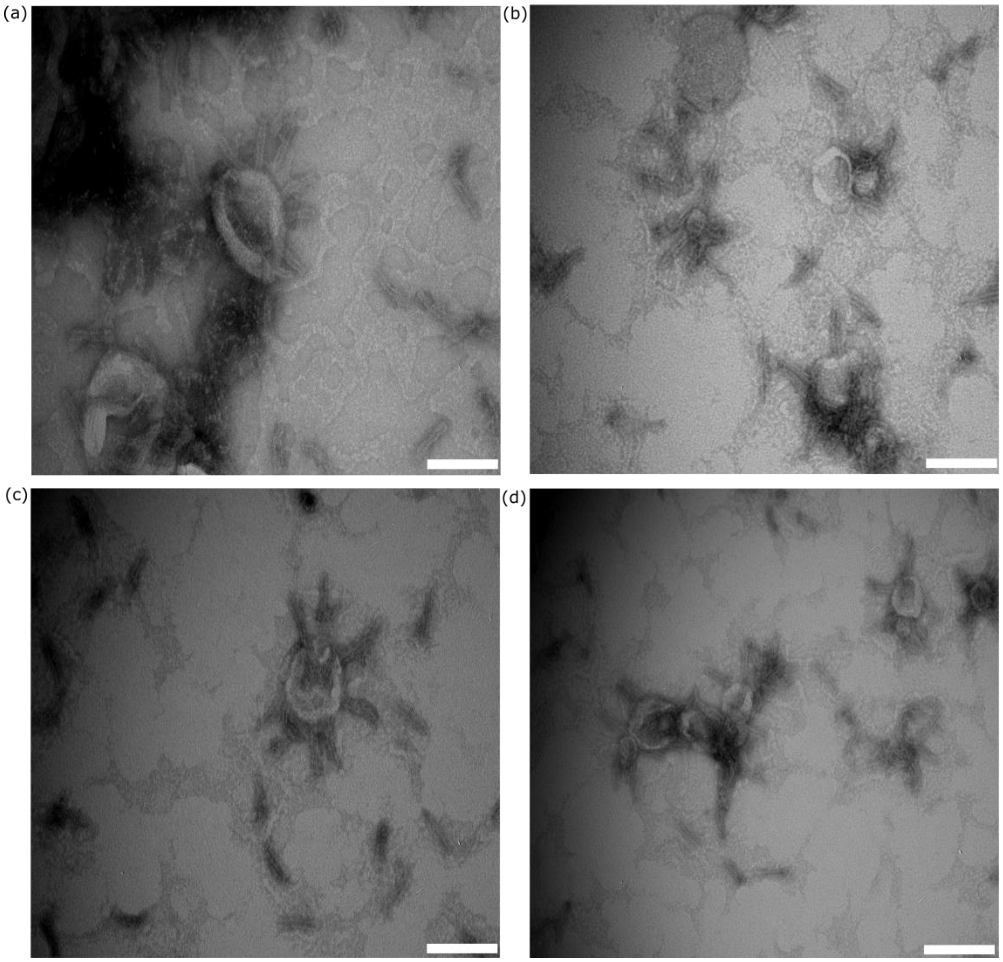


**Figure S5.** (**a–d**) TEM micrographs of DNA origami channels combined with small unilamellar vesicles (SUVs). DNA origami seeds penetrate the SUVs. Some of the seeds are also aggregated inside the SUVs, possibly due to breakage of the SUVs on the TEM grids. Scale bars: 100 nm (see Materials and Methods for protocol).

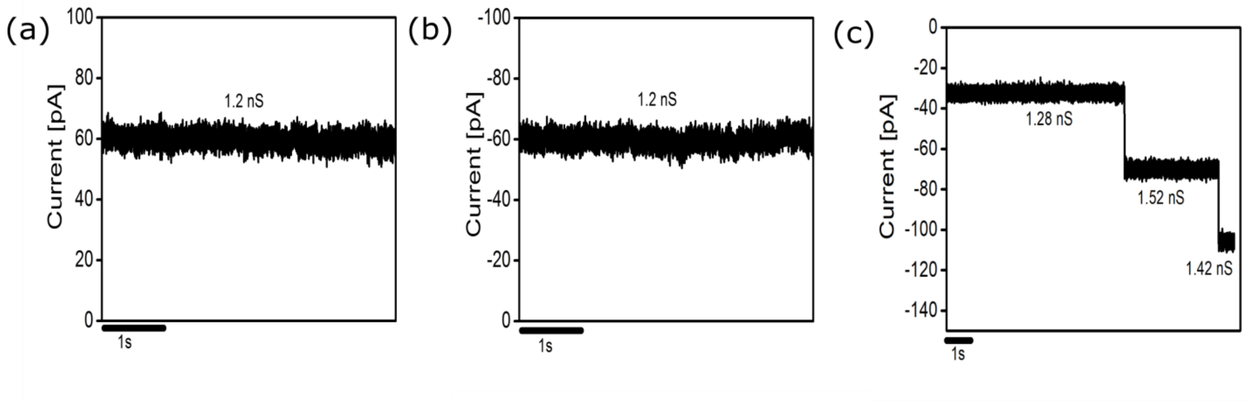


**Figure S6.** Typical ion-current traces showing insertions of the DNA origami nanotube seed channels at applied transmembrane potentials of (**a**) +50 mV; (**b**) –50 mV; and (**c**) –25 mV. The conductances corresponding to each insertion are labeled on the graphs. The electrical measurements were carried out in 1 M KCl, 10 mM $MgCl_2$ buffered with 10 mM 4-morpholineethanesulfonic acid (MES) monohydrate, pH 6.0. For clarity, the traces were filtered at 200 Hz.

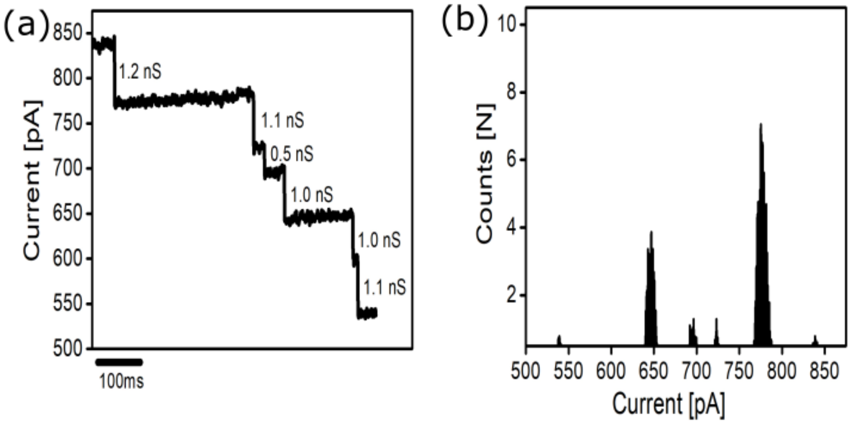


**Figure S7.** (**a**) Electrical traces showing the closure of the DNA origami nanotube seed channels. The total conductance of all nanotube seeds in the given trace is 5.9 nS. Here the two droplets are separated from each other resulting in the separation of the lipid bilayer into a monolayer. While separating the droplets, 6 single-channel closures occurred at an applied transmembrane potential of +50 mV; and (**b**) The corresponding all-points histogram of the 6 single-channel closures showing the variation in conductances of different structures and over time. The electrical measurements were carried out in 1 M KCl, 10 mM $MgCl_2$ buffered with 10 mM MES, pH 6.0. For clarity, the traces were filtered at 200 Hz.

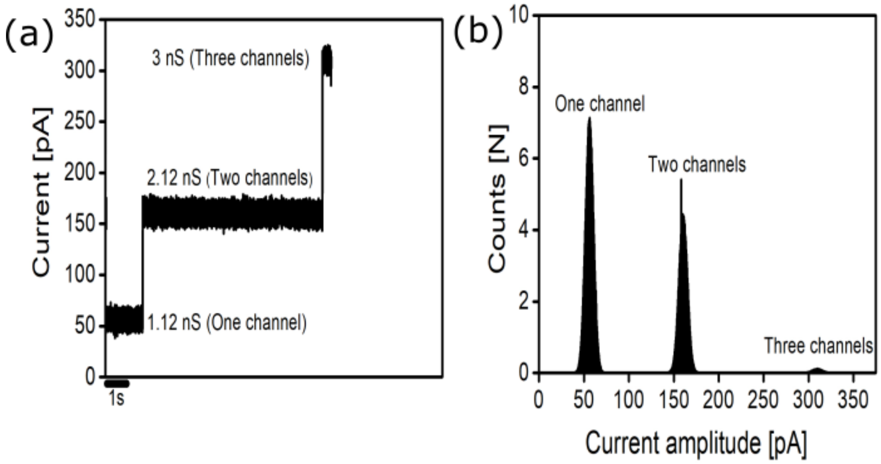


**Figure S8.** (**a**) Electrical trace consistent with the insertion of one DNA origami nanotube seed channel, followed by the simultaneous insertion of two DNA origami seeds channels followed by the simultaneous insertion of three DNA origami seeds channels. Measurements were made at an applied transmembrane potential of +50 mV; and (**b**) The corresponding all-points histogram of (**a**). The electrical measurements were carried out in 1 M KCl, 10 mM $MgCl_2$ buffered with 10 mM MES, pH 6.0.

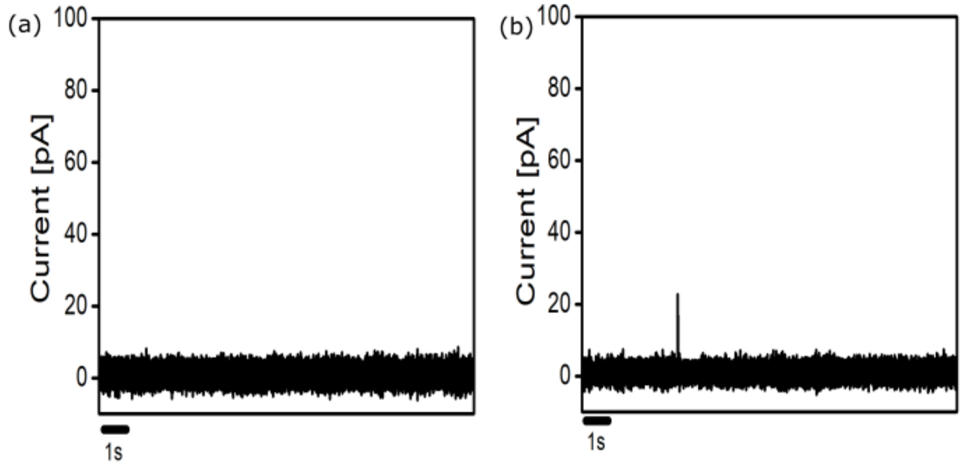


**Figure S9.** Control experiments for DNA origami nanotube seed channels. (**a**) Sample ion-current trace of a solution of DNA origami nanotube seeds to which no DNA-cholesterol conjugate strand (3' nanotube seed_Chol.DNA) were attached at an applied transmembrane potential of +50 mV. Traces recorded showed no signs of conductance pattern of more than 10 pS recorded for about 30 min in 1,2-diphytanoyl-*sn*-glycero-3-phosphocholine (DPhPC) bilayers. This suggests that there is no channel insertion; and (**b**) Sample ion-current trace of 10 µM DNA-cholesterol (3' nanotube seed_Chol.DNA) conjugate at an applied transmembrane potential of +50 mV. Traces recorded showed no signs of conductance pattern of more than 10 pS recorded for about 30 min in DPhPC bilayers. This suggest that there is no channel insertion. The electrical measurements were carried out in 1 M KCl, 10 mM $MgCl_2$ buffered with 10 mM MES, pH 6.0.

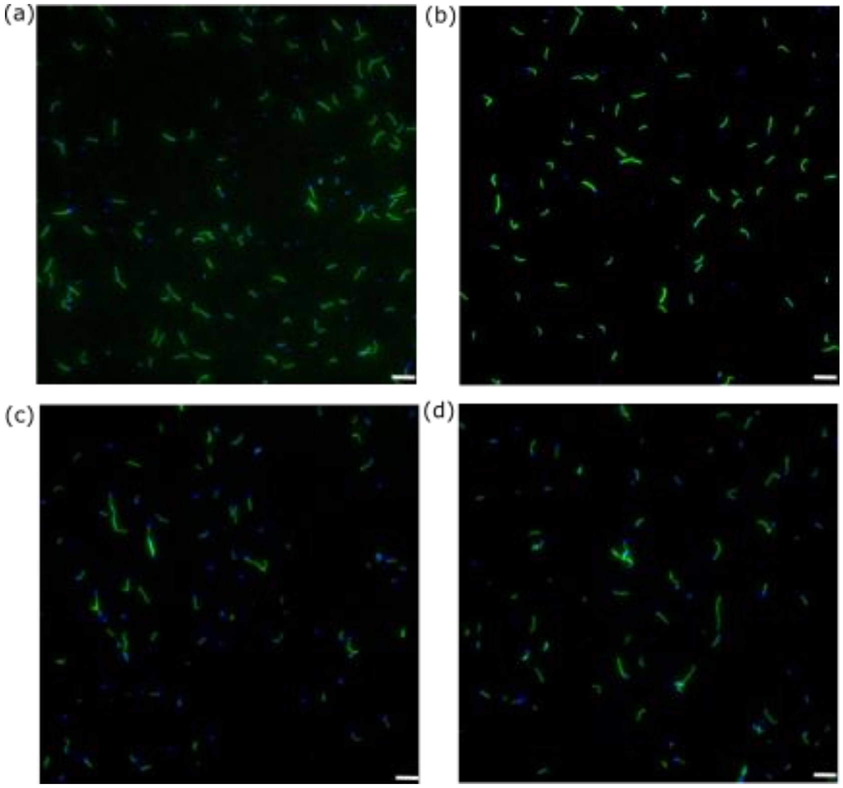


**Figure S10.** Fluorescence micrographs of seeded DNA nanotubes. (**a**, **b**) Seeded nanotubes before filter purification; (**c**, **d**) Seeded nanotubes after 3 rounds of filter purification at 3000 g for 30 min. The nanotube seeds were labeled with (ATTO488, blue) and the nanotubes were labeled with (Cy3, green). Scale bars: 5 µm. The detailed annealing and purification protocol is given in Notes S6 and S7.

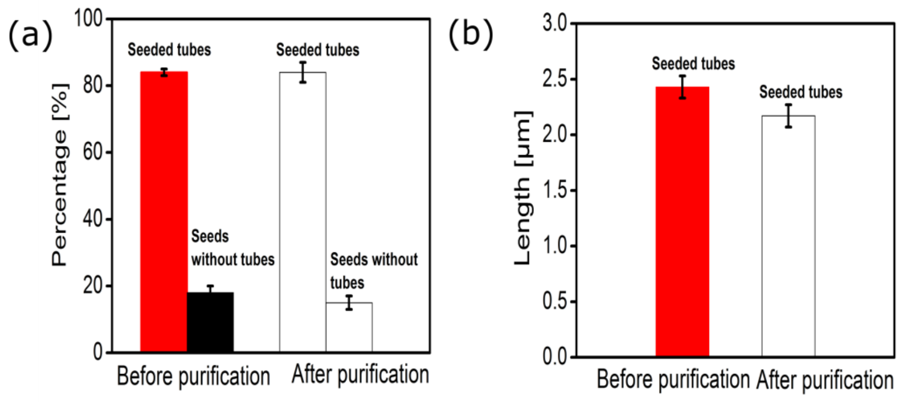


**Figure S11.** DNA seeded nanotubes (**a**) Before three rounds of filter purification in 1X TAEM buffer yields 84 ± 1% (95% CI) (N = 467 structures) of tubes with seeds. After three rounds of filter purification, 84 ± 3% (95% CI) (N = 678 structures) of seeds had attached nanotubes; and (**b)** Length of the DNA nanotubes before and after filter purification. Before filter purification the average length of seeded nanotubes was 2.43 ± 0.10 µm (95% CI) (N = 185) and after filter purification the average length was 2.17 ± 0.10 µm (95% CI) (N = 272).

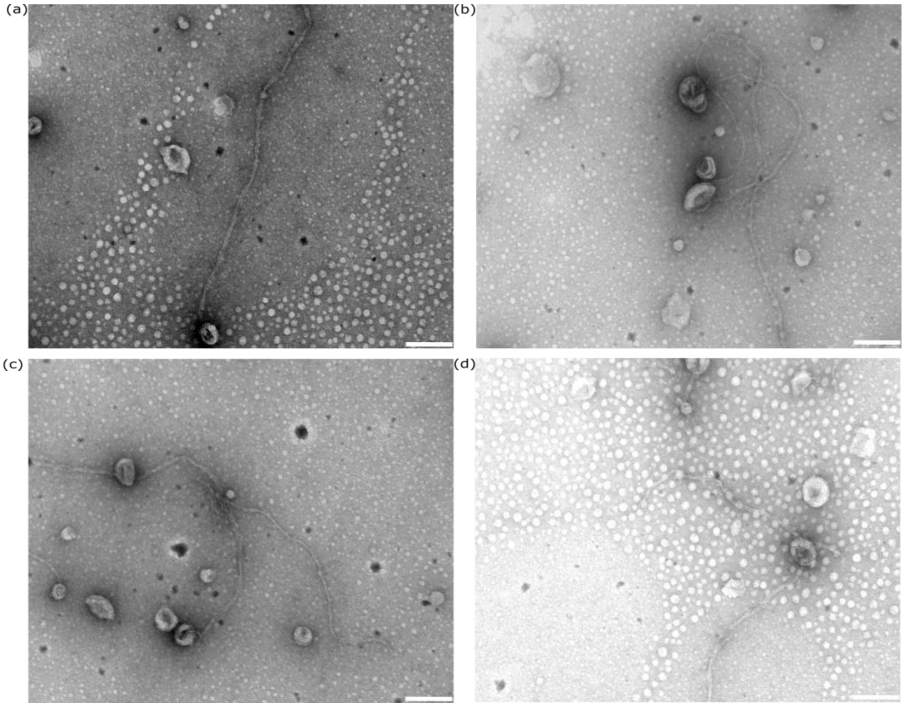


**Figure S12.** (**a–d**) TEM micrographs showing the incorporation of DNA nanotubes channels into SUVs. The DNA nanotubes that insert are at least 1 micron in length. Scale bars: 100 nm (see Materials and Methods for protocol).

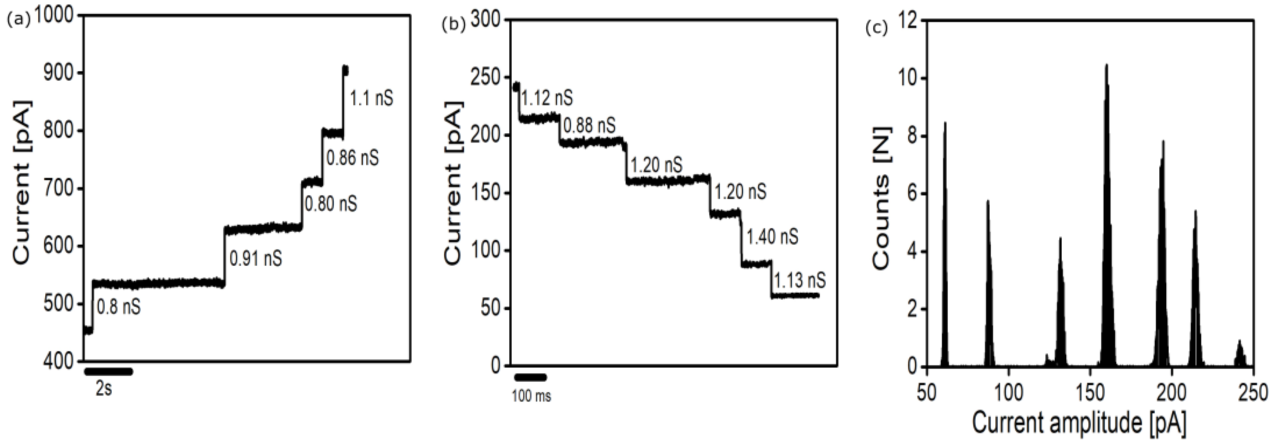


**Figure S13.** Electrical signatures of the DNA seeded nanotube channel insertions and closure/departure from DPhPC bilayers during droplet separation. (**a**) Multiple insertions of the seeded nanotube channels at an applied transmembrane potential of +100 mV. The total conductance change from the ion current jumps shown is 4.47 nS; (**b**) Six drops in current that occurred while separating the two droplets forming a bilayer at an applied transmembrane potential of +25 mV; and (**c**) The corresponding all-points histogram of (**b**). The electrical measurements were carried out in 1 M KCl, 10 mM $MgCl_2$ buffered with 10 mM MES, pH 6.0. For clarity, the traces were filtered at 200 Hz.

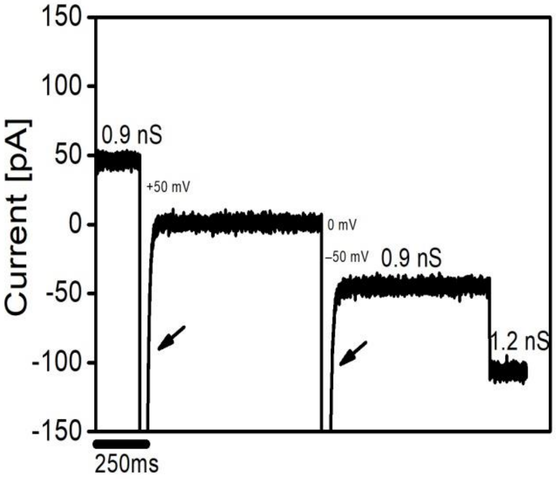


**Figure S14.** Additional multi-channel insertions of the DNA seeded nanotube channels at both +50 mV and –50 mV respectively. The black arrow indicates the change of transmembrane potential. The electrical measurements were carried out in 1 M KCl, 10 mM $MgCl_2$ buffered with 10 mM MES, pH 6.0. For clarity, the traces were filtered at 200 Hz.

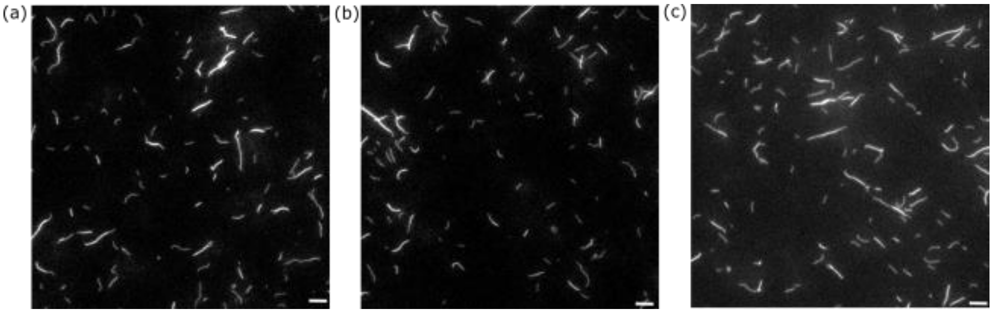


**Figure S15.** Fluorescence micrographs of unseeded DNA nanotubes grown at 32°C for >18 h. The samples were diluted 50 times prior to imaging them on an inverted microscope (Olympus IX171) using a 60x/1.45 NA oil immersion objective using an Olympus Cy3 filter cube set (Z532BP). Scale bars: 5 µm. The detailed annealing protocol is given in Note S8.

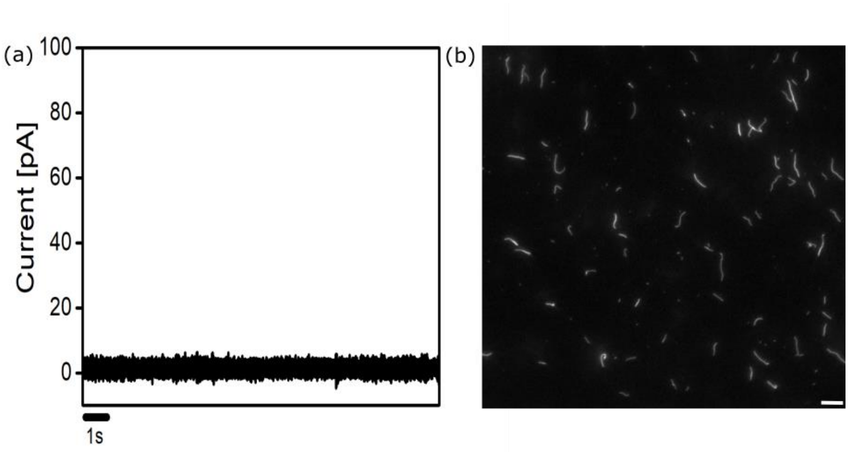


**Figure S16.** (**a**) Ion-current trace of unseeded DNA nanotubes in DPhPC bilayers. Nanotubes were prepared by annealing 250 nM each REd and SEd monomers to 32°C and then incubating them for >18 h. Annealing and purification of the unseeded nanotubes are performed as described in Note S8. Traces recorded over 30 min showed no signs of conductance pattern of more than 10 pS in DPhPC bilayers. This suggest that there is no channel insertion. The electrical measurements were carried out in 1 M KCl, 10 mM $MgCl_2$ buffered with 10 mM MES, pH 6.0 at an applied transmembrane potential of +50 mV; and (**b**) Fluorescence micrographs of the unseeded nanotubes after annealing and filter purification (see Note S8). Nanotube samples were imaged them on an inverted microscope (Olympus IX171) using a 60x/1.45 NA oil immersion objective using an Olympus Cy3 filter cube set (Z532BP). Scale bar: 5 µm.

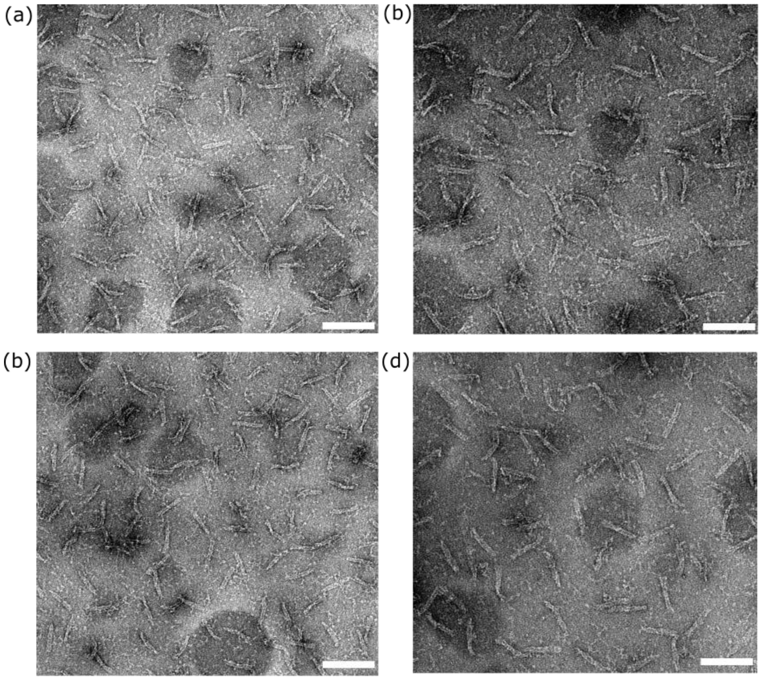


**Figure S17.** TEM micrographs of DNA origami channel caps after annealing and filter purification as described in Note S9. Scale bars: 100 nm.

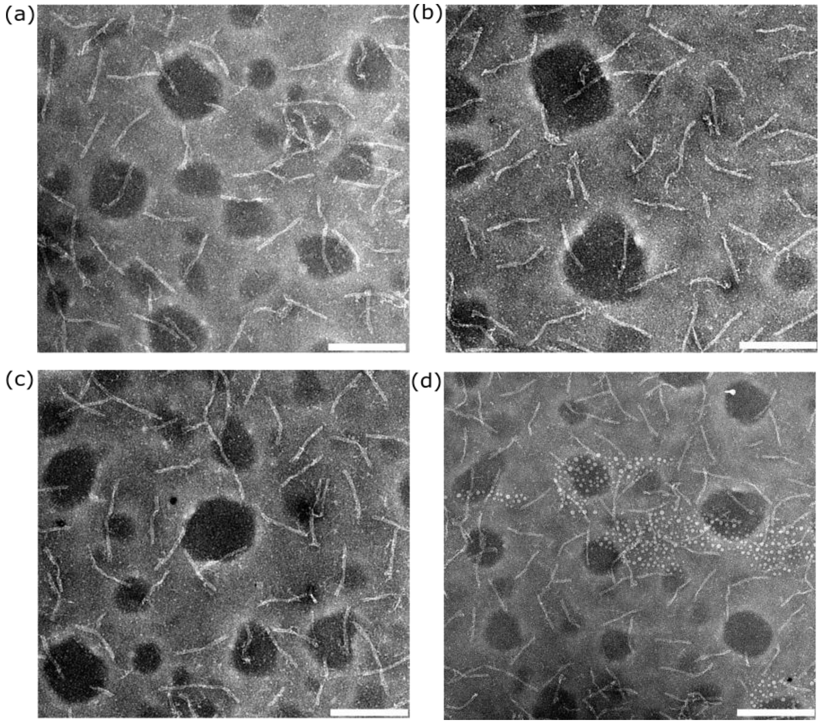


**Figure S18.** (**a–d**) Transmission electron micrographs of DNA nanotube seeds bound to channel caps. The complexes were prepared and purified as described in Note S9. Scale bars: (**a**) and (**b**)100 nm; (**c**) and (**d**) 200 nm.

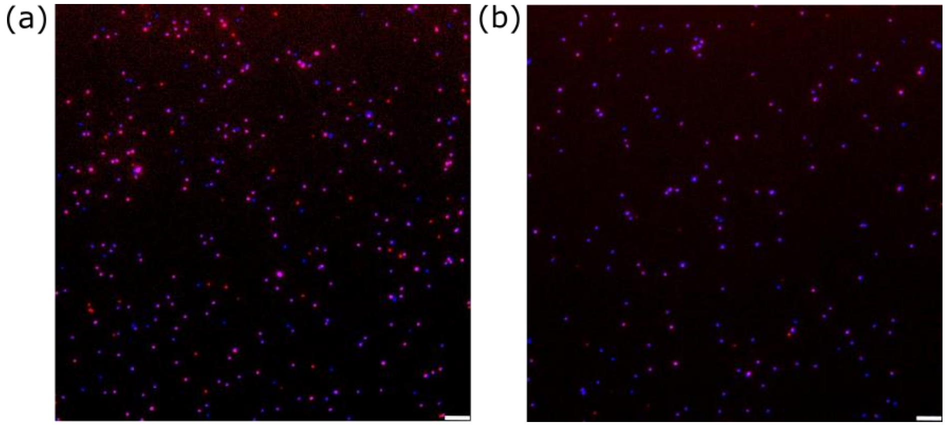


**Figure S19.** Fluorescence micrographs of DNA nanotube seeds bound to channel caps. The seed-channel cap complexes were prepared as described in Note S9. (**a**) Seed channel cap complexes after annealing but before filter purification; and (**b**) Seed-channel cap complexes after 3 rounds of filter purification at 3000 g for 30 min per round. There was no significant change in concentration of channel capped nanotube seeds before and after 2 rounds of filter purification as observed from Figure S20. Scale bars: 5 µm. The nanotube seeds were labeled with ATTO488, blue and the channel caps with ATTO647N, red.

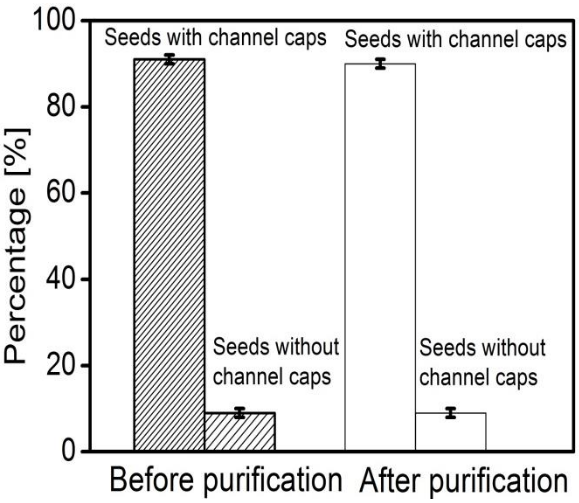


**Figure S20.** Percentages of DNA origami nanotube seeds that were bound *vs.* not bound to channel caps. Before filter purification, 91 ± 1% (95% CI, N = 230) of seeds had attached channel caps. After filter purification containing the cholesterol-DNA conjugate (3' nanotube seed_Chol.DNA) 90 ± 1% (95% CI, N = 223) of nanotube seeds still bound to channel caps. Filter purification was carried in 1X TAEM buffer at 3000 g for 30 min 1 time. The detailed annealing and purification protocol is given in Note S9.

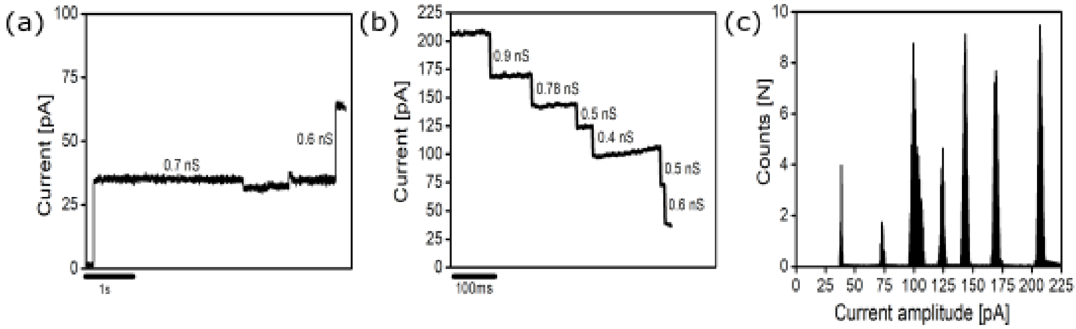


**Figure S21.** Typical multi-channel insertional trace of the nanotube seeds with channel caps in 1 M KCl, 10 mM $MgCl_2$ buffered with 10 mM MES, pH 6.0 at an applied transmembrane potential of (**a**) +50 mV; (**b**) The separation of the two droplets results in the monolayer formation which leads to the closure of the channel capped origami seed channels leading to a total conductance of 3.68 nS; and (**c**) The corresponding all-points histogram of (**b**).

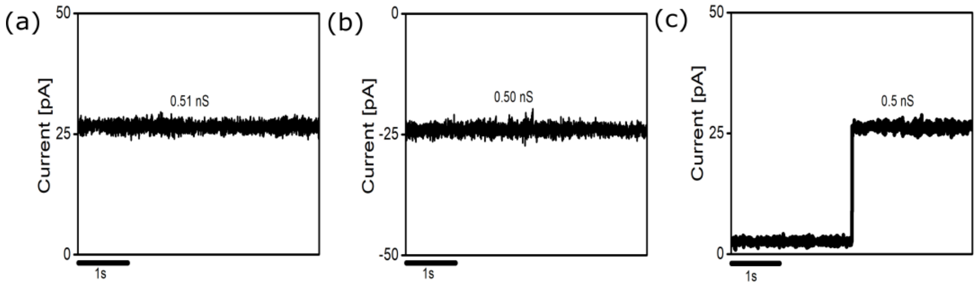


**Figure S22.** Additional single-channel electrical signatures of single nanotube seeds with channel caps in 1 M KCl, 10 mM $MgCl_2$ buffered with 10 mM MES, pH 6.0 at an applied transmembrane potential of (**a** and **c**) +50 mV; and (**b**) –50 mV respectively.

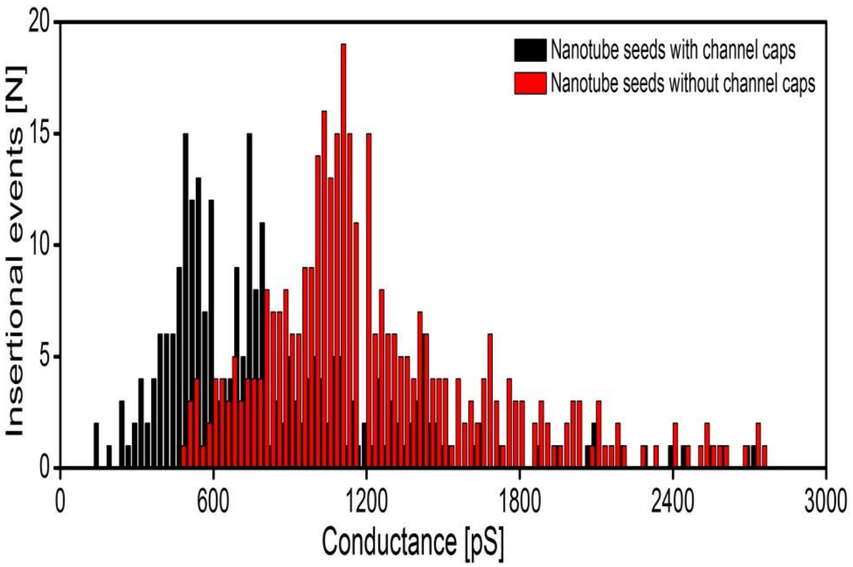


**Figure S23.** The single-channel conductance histograms from main text **Figures 2h** and **4h** overlayed to make it possible to compare the open channel insertion of the nanotube seeds and nanotube seeds with channel caps. The conductances of both structures were measured in a buffer consisting of 1M KCl, 10 mM $MgCl_2$ buffered with 10 mM MES, pH 6.0 at an applied transmembrane potential of +50 mV.

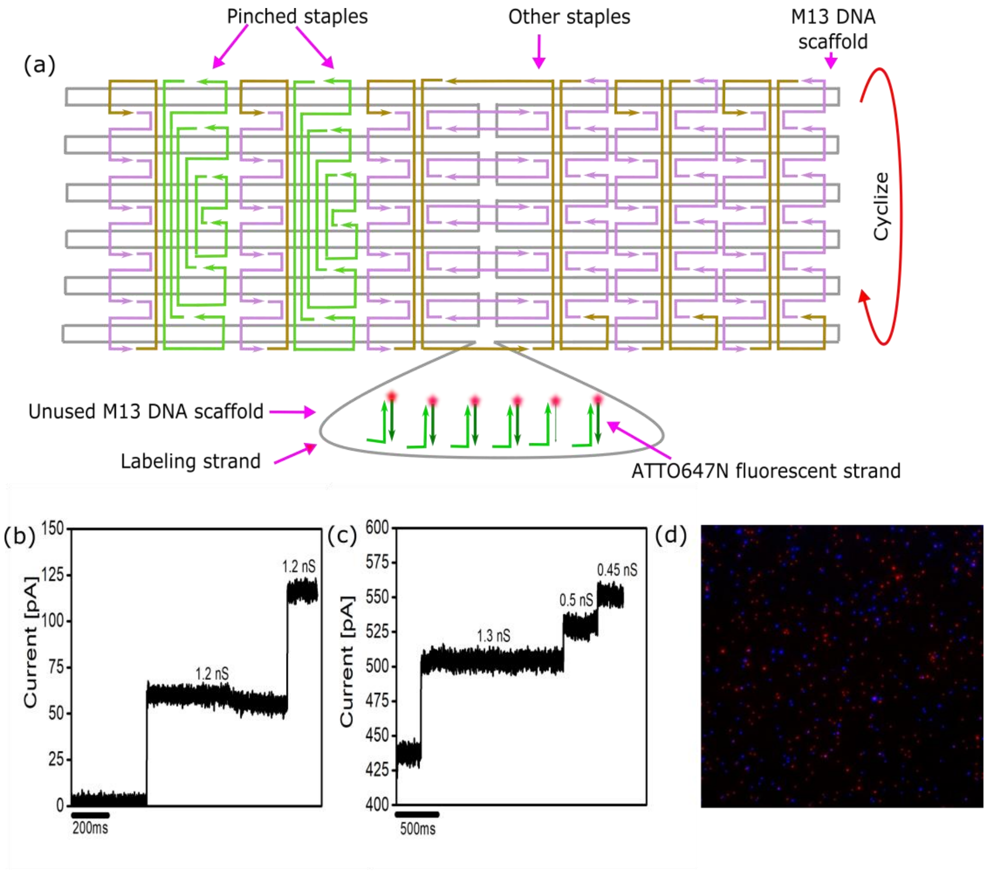


**Figure S24.** (**a**) Schematic showing the modified Cadnano map[1–3]of the staples and adapters of the control DNA origami channel cap. The 6 adapter strands presenting the sticky ends that can bind to a seed or nanotube end have been removed from the structure; (**b** and **c**) Ion-current signatures of the DNA nanotube seed channels incubated with control channel caps as described in Note S11; and (**d**) Multi-color fluorescence micrographs of the DNA nanotube seed channels and channel caps after the two species were incubated together as described in Note S11. DNA nanotube seed channels did not attach to the control channel caps. DNA nanotube seeds are labeled with ATTO488 (blue) and channel caps with ATTON647 (red). Scale bar: 5 µm.

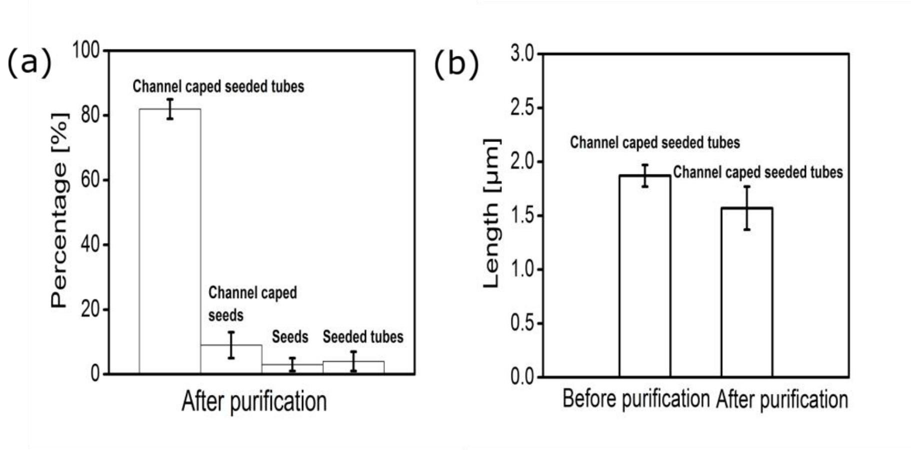


**Figure S25.** Channel capped nanotubes (**a**) after two rounds of filter purification in 1X TAEM buffer yields 82 ± 3% (95% CI) of capped seeded tubes, 9 ± 4% (95% CI) of capped seeds, 3 ± 2 (95% CI) and 4 ± 3 (95% CI) of the seeds and seeded tubes alone in the solution; and (**b**) Length of the channel capped DNA nanotubes before and after filter purification at 3000 g for 30 min, performed twice. Before filter purification, the average length of seeded nanotubes was 1.87 ± 0.10 µm (95% CI, N = 108) and after filter purification the average length was 1.57 ± 0.20 µm (95% CI, N = 105), respectively. The annealing and purification protocol is given in Note S10.

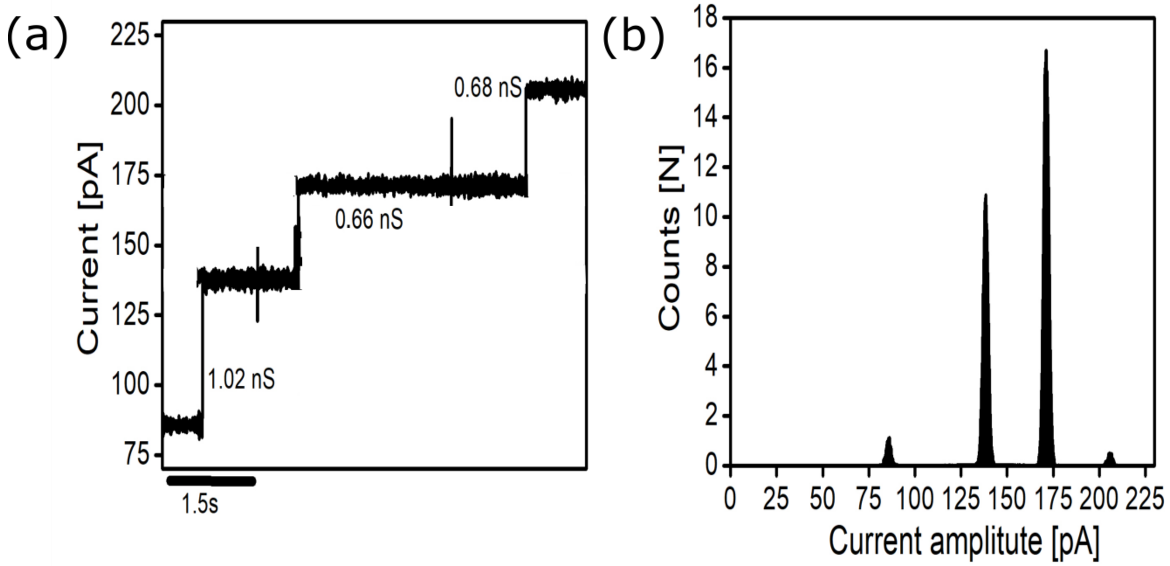


**Figure S26.** Typical multi-channel insertional trace of the channel capped seeded nanotubes in 1 M KCl, 10 mM $MgCl_2$ buffered with 10 mM MES, pH 6.0 at an applied transmembrane potential of (**a**) +50 mV; and (**b**) The corresponding all-points histogram of (**a**).

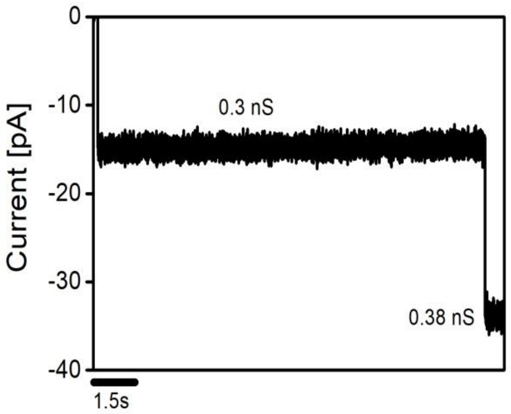


**Figure S27.** Additional electrical signature of a channel capped seeded nanotube in 1 M KCl, 10 mM $MgCl_2$ buffered with 10 mM MES, pH 6.0 at an applied transmembrane potential of –50 mV.

### References


1 S. M. Douglas, A. H. Marblestone, S. Teerapittayanon, A. Vazquez, G. M. Church, W. M. Shih, *Nucleic. Acids. Res.*, 2009, **37**, 5001–5006.

2 S. M. Douglas, H. Dietz, T. Liedl, B. Högberg, F. Graf, W. M. Shih, *Nature*, 2009, **459**, 414–418.

3 H. Dietz, S. M. Douglas, W. M. Shih, *Science*, 2009, **325**, 725–730.

4 A. M. Mohammed, P. Šulc, J. Zenk, R. Schulman, *Nat. Nanotechnol.*, 2017, **12**, 312–316.

5 A. M. Mohammed, R. Schulman, *Nano Lett.*, 2013, **13**, 4006−4013.

6 D. K. Agrawal, R. Jiang, S. Reinhart, A. M. Mohammed, T. D. Jorgenson, R. Schulman, *ACS Nano,* 2017, **11**, 9770–9779.

7 M. Langecker, V. Arnaut, T. G. Martin, J. List, S. Renner, M. Mayer, H. Dietz, F. C. Simmel, *Science,* 2012, **338**, 932–936.
